# Structure-guided assembly of an influenza spike nanobicelle vaccine provides pan H1 intranasal protection

**DOI:** 10.1101/2024.09.16.613335

**Authors:** Mallory L. Myers, John R. Gallagher, De’Marcus D. Woolfork, Noah D. Khorrami, William B. Park, Samantha Maldonado-Puga, Eric Bohrnsen, Benjamin H. Schwarz, Derron A. Alves, Kevin W. Bock, Altaira D. Dearborn, Audray K. Harris

**Affiliations:** Structural Informatics Unit, Laboratory of Infectious Diseases, National Institute of Allergy and Infectious Diseases, National Institutes of Health, 50 South Drive, Room 6351, Bethesda, MD, USA 20892; Protein Chemistry Section, Research and Technologies Branch, National Institute of Allergy and Infectious Diseases, Rocky Mountain Laboratories, National Institutes of Health, 903 South 4th Street, Hamilton, MT, USA 59840; Infectious Disease Pathogenesis Section, National Institute of Allergy and Infectious Diseases, National Institutes of Health, 33 North Drive, Room BN25, Bethesda, MD, USA 20892; Structural Virology Section, Laboratory of Infectious Diseases, National Institute of Allergy and Infectious Diseases, National Institutes of Health, 50 South Drive, Room 6531, Bethesda, MD, USA 20892

**Author notes:** To whom correspondence should be addressed; E-mail A. K. H.

## Abstract

Development of intranasal vaccines for respiratory viruses has gained popularity. However, currently only a live-attenuated influenza vaccine is FDA-approved for intranasal administration. Here, we focused on influenza virus as it circulates seasonally, has pandemic potential, and has vaccine formulations that present hemagglutinin (HA) in different structural arrangements. These display differences have not been correlated with induction of pan-H1 antibodies or shown to provide intranasal protection. Using electron microscopy, biochemistry and animal studies, we identified HA complexes arranged as lipid discs with multiple trimeric HAs displayed along the perimeter, termed spike nanobicelles (SNB). We utilized a structure-guided approach to synthesize in vitro assembled spiked nanobicelles (IA-SNB) from a classical 1934 H1N1 influenza virus. IA-SNBs elicited pan-H1 antibodies and provided protection against antigenically divergent H1N1 viruses via intranasal immunizations. Viral glycoprotein spikes displayed as SNBs could aid in combating antigenic variation and provide innovative intranasal vaccines to aid universal influenza vaccine development.

## Introduction

Influenza viruses infect millions of people worldwide on an annual basis (seasonal influenza) and influenza can cause global pandemics like the 1918 and 2009 pandemics caused by H1N1 influenza viruses^1,2^. Commercial vaccines target the major surface glycoprotein hemagglutinin (HA), but constant antigenic variation of HA requires vaccines to be reformulated every year^3–5^. Antigenic change in HA can be described by two phenomena: antigenic drift and antigenic shift^6^. Rapid antigenic drift powers a constantly renewed ability to evade the host immune response, making influenza virus a significant challenge for public health^3–5^. Antigenic shift, facilitated by the segmented influenza viral genome structure, can give rise to reassortment between animal and human influenza viruses, leading to the emergence of antigenically novel influenza viruses and causing pandemics that pose significant risks to even healthy populations^7–9^. As a result, the production of seasonal influenza vaccines has become a billion-dollar industry^10^.

Influenza is caused by infection with an enveloped virus which has three surface glycoproteins: hemagglutinin (HA) and neuraminidase (NA) and Matrix (M2). The vast majority of protective antibodies following natural influenza virus infection target HA^11–13^, a 3-fold symmetric spike^14^. Influenza vaccines contain H1, H3 type A and type B influenza strains, reflecting the most common influenza subtypes circulating in humans^4,15,16^. Commercial influenza vaccines have been available in the United States since the 1950s^17^, with the Center for Disease Control (CDC) regulating both the strain of H1, H3, and influenza B HA components as well as the dose of HA formulated in the vaccines^11^. In 2015, the Food and Drug Administration (FDA) approved Fluad, the first influenza vaccine containing an adjuvant, MF59^18^.

Although structure-guided designs of symmetrical nanoparticles displaying HA ectodomain fragments have been evaluated in detail^19–22^, the structure-guided evaluation and formulation of full-length HA vaccines such as those implemented in commercial influenza vaccines have not received a similar level of study. Differences in the structural arrangements of constituent HA molecules in various commercial influenza vaccine formulations have been shown, including the number of HAs per molecular complex, as well as the structural relationship of HAs within these complexes^23^. HA complexes in current commercial influenza vaccines generally fall into four main configurations: (i) isolated HA molecules, (ii) starfish-like complexes, (iii) HA in micelles and (iv) large disc-like structures^23–26^. HA was only seen in large disc-like structures in the adjuvanted vaccine, Fluad, and this correlated with the elicitation of higher levels of heterosubtypic cross-reactive antibodies^23^.

However, to date most structure-guided design efforts to enhance and to target antibody responses have focused on designing symmetrical nanoparticle systems. Examples, include the use of an octahedral ferritin particle as an effective nanoparticle scaffold to display HA ectodomains ^20–22,27–29^. However, these ferritin-based particles used adjuvant via intramuscular immunization to elicit protective immune responses^30–32^. In contrast intranasal influenza vaccination for mucosal immunity targets the immune response to the route of infection, but only live-attenuated influenza viruses have been FDA approved (i.e. Flumist) for intranasal immunization^33,34^. One problem is that antigen instability can affect the efficacy of influenza vaccines^35,36^. There are concerns for eliciting adverse reactions from intranasal adjuvanted vaccines such as possible neurological disorders^37^. Thus, there a need for intranasal vaccine development that use other platforms for HA display to the immune system besides live-attenuated viruses and adjuvanted subunit vaccines.

In this work, we explore the effects of adjuvant removal and of structural organizations of HA complexes in the form of “spiked nanobicelles” (SNB) for eliciting intranasal pan-H1 protection. We found that SNBs could be purified from the Fluad vaccine. Lipidomic analysis indicated an enrichment in long chain sphingolipids suggestive of a stable, high melting point, lipid architecture to SNB discs. We then developed a protocol for the in vitro assembly of SNBs from the classical A/Puerto Rico/8/34 (H1N1) virus produced from eggs. We termed these H1 1934 complexes as in vitro assembled spiked nanobicelles (IA-SNB). The H1 1934 IA-SNBs were determined to be structurally and compositionally similar to Fluad SNB by electron microscopy and lipidomics. 1934 IA-SNBs provided pan-H1N1 protection to mice via intranasal immunizations. These protections were from lethal challenges with antigenically matched (1934 H1N1) and antigenically mismatched pandemic (2009 H1N1) and seasonal (2015 H1N1) influenza viruses. The pan-H1 breadth of cross-reactivity and protection included influenza H1 subtypes strains spanning over 80 years. These results herald an important step forward in the structure-guided design of intranasal vaccines for the elicitation of broadly protective immune responses against antigenically variable influenza viruses. In addition, spike nanobicelles could not only aid efforts to development universal influenza vaccines but also aid intranasal vaccine development for other respiratory viruses.

## Results

### Electron microscopy and characterization of Fluad

To further understand the organization of spiked nanobicelles (SNBs), Fluad was imaged by cryo-electron microscopy (cryo-EM) and SNBs were biochemically purified and subjected to lipidomic analysis. Fluad, a commercially available influenza vaccine, is manufactured from formaldehyde-inactivated split virus mixed with MF59 adjuvant (Supplementary Fig. 1). Cryo-electron microscopy (cryo-EM) indicated that there were two major structural components of Fluad (Fig. 1a), spiked nanobicelles (SNB) (Fig. 1a, white arrows) and particles of MF59 adjuvant (Fig. 1a, black arrows). The MF59 adjuvant was the more abundant structural component. Adjuvant particles had an average diameter of 88 nm (n=84) and ranged from 38 nm to 163 nm in diameter. The density inside the adjuvant appeared to be higher (darker gray) than the surrounding buffer (lighter gray) (Fig. 1a), indicating that adjuvant particles are not empty vesicles but micelles with hydrophobic cores. A schematic 3D model of the Fluad vaccine displays adjuvant as spherical particles with interspersed spiked nanobicelles (Fig. 1b). SNBs are lipid discs surrounded by two perimeter rings of hemagglutinin (HA) molecules on top and bottom. This SNB structure is not detected in other commercial influenza vaccines which contained either smaller HA-complexes (i.e. Flublok) or larger influenza virions (Flumist) when examined by electron microscopy (Supplementary Fig. 2).

**Fig. 1.**
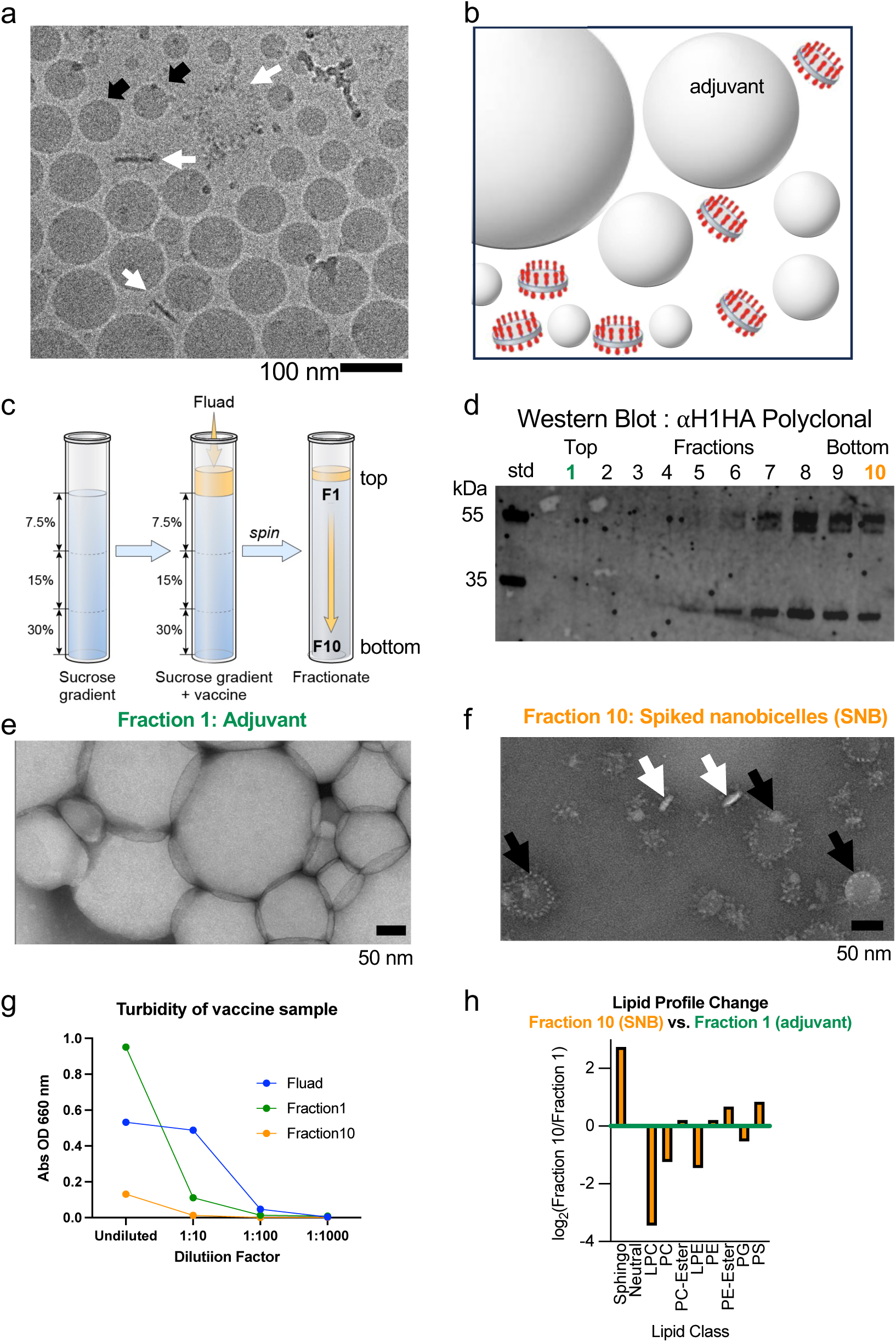
Characterization of MF59 adjuvant particles and spiked nanobicelles particles by electron microscopy and lipidomics. (a) cryo-electron microscopic (EM) image of Fluad with adjuvant particles (black arrows) and spiked nanobicelles (SNB) (white arrows). Scale bar 100 nm. (b) Schematic of spiked nanobicelles with hemagglutinin (HA) (red) and membrane (grey) dispersed among adjuvant particles. (c) Schematic of gradient used to separate adjuvant and purify spiked nanobicelles (d) Western blot of fractions taken from the top to the bottom of the gradient to probe for the present of HA protein via anti-H1 HA polyclonal antibody. (e, f) Negative-staining EM image of top fraction #1 showing adjuvant particles (white vesicles) (panel e) and EM image of bottom fraction #10 showing a field of purified spiked nanobicelles (panel f). White arrows indicate approximate side views of the spike nanobicelles with the membrane being white ovals. Black arrows indicate approximate top views of the spike nanobicelles with a roughly circular shape with HA spikes on the perimeter the membrane being white ovals. Scale bars 50 nm. (g) Turbidity measurements of fraction 1, fraction 10 and Fluad (control) to monitor the separation of adjuvant from spiked nanobicelles. (h) Lipid profile with lipid classes that increase (positive values) and decrease (negative values) for fraction 10 (spiked nanobicelles) when compared to fraction 1 (adjuvant).

### Separation of Fluad components: spiked nanobicelles and MF59 adjuvant particles

Fluad was biochemically separated using gradient centrifugation, sucrose step gradient (7.5-30%) (Fig 1c). Prior to centrifugation, the Fluad vaccine visually appeared as a milky white band on top of the gradient (Supplementary Fig. 3a). Following centrifugation, the milky band decreased in thickness but remains on the top of the gradient (Supplementary Fig. 3b). Fractions collected from the top to the bottom of the gradient (fractions 1 to 10) varied in HA content, with HA detected only in the bottom half of the gradient (fractions 5-10) by immunoblot (Fig. 1d). Western blot analysis indicated that fraction 10 was dominated by HA (Fig 1d) and other viral components, such as neuraminidase (NA), nucleoprotein (NP), and matrix (M1) were not detectable (Supplementary Fig. 4). Additionally, the higher sucrose density of fraction 10 should have macromolecular complexes of megadalton sizes (i.e., spiked nanobicelles (SNBs) of HA).

When fraction 1 was examined by negative-staining electron microscopy there is detection of large white particles indicating that fraction 1 contained MF59 adjuvant particles (Fig. 1e). In contrast, fraction 10 contained flat discs displaying HA spikes, SNBs (Fig. 3f). Some SNB were oriented as side views with white flat discs of membrane with protruding HA spikes (Fig. 3f, white arrows), while top views appeared as white discs of membrane ringed by perimeters of protruding HA spikes (Fig. 3f, black arrows). There was no indication of MF59 adjuvant like structures detected in these purified SNBs (fraction 10) as judged by electron microscopy (Fig. 1e, versus Fig. 1f).

### Relative turbidity and lipidomic classification of MF59 adjuvant particles and spiked nanobicelles

The turbidities of the fractionated samples (fraction 1, green. fraction 10, orange) were compared to the turbidity of Fluad (non-fractionated, blue). Fraction 1 (adjuvant) had highest and fraction 10 the lowest turbidities (Fig. 1g). Following a 1:10 dilution, the Fluad control maintained a similar level of turbidity to the undiluted reading while both spiked nanobicelles (SNB) (fraction 10) and adjuvant (fraction 1) samples had lower levels of turbidity. When the dilution factor was increased 100-fold there was no measurable turbidity for any of the samples (Fig. 1g). Lipidomic analysis demonstrated that SNB (fraction 10) was enriched greater than 8-fold for sphingolipids compared to the lower density Fluad 1 fraction, in particular ceramides and hexosylceramides with either saturated or very long chain (>C20) acyl-groups (Fig. 1h) (Supplementary Fig. 5a).

### Structural analysis of spiked nanobicelles by cryo-electron microscopy

In order to study the structure of spiked nanobicelles (SNBs) and the conformation of the constituent HA molecules cryo-EM image analysis and 3D reconstruction (∼4.1 Å resolution) were performed on purified spiked nanobicelles (fraction 10) (Supplementary Figs. 6, 7, 8, 9). Side-views of the single images of spiked nanobicelles (Fig. 2a) and 2D class averages (Fig. 2b, 2c) indicated that SNBs consisted of a lipid bilayer (∼10.6 nm thick) with about 4.3 nm of spacing between the bilayers consistent with the very long chain ceramides observed in the lipidomics data^38^. There were top and bottom layers of rings of HA molecules on the perimeter of the bilayer with HA molecules embedded in the membrane via transmembrane regions (Fig. 2a-2e). When measured, SNBs in the side-view orientations (n=94), had an average height of 37 nm (range 29 nm to 53 nm) and an average diameter of 42 nm (range 24 nm to 68 nm) (Fig. 2a). These were smaller than those SNBs observed in the top-view orientations (n=65) that had an average diameter of 90 nm (range 55 nm to 130 nm) (Fig. 2d). This suggests that SNBs of different diameters might have preferred orientations in cryo-EM.

**Fig. 2.**
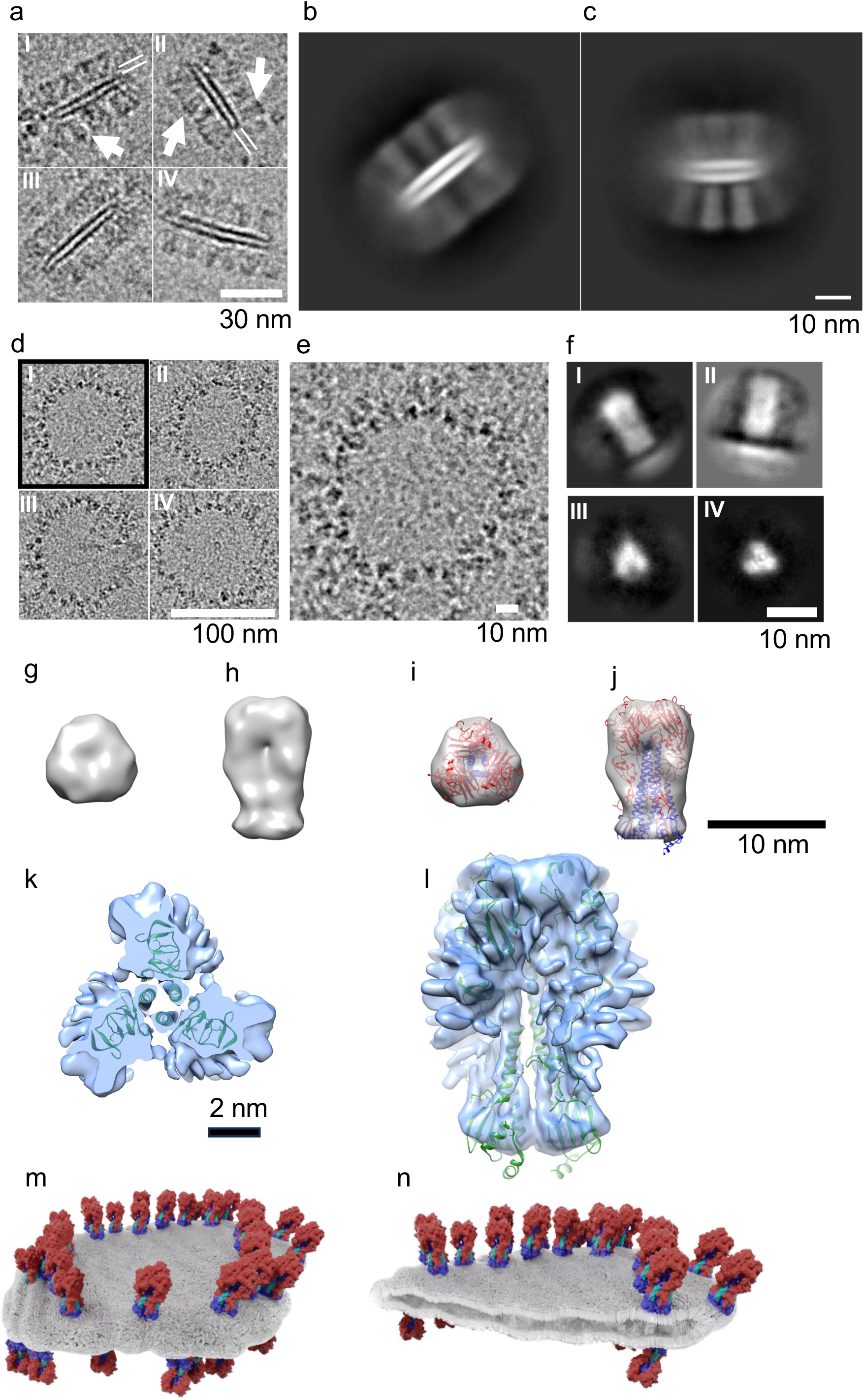
Organization of spiked nanobicelles by cryo-electron microscopy (cryo-EM) and image analysis. (a) Montage of cryo-EM images (side-views) of spiked nanobicelles (I-IV). For panels I, II, spikes of HA molecules are denoted (white arrows) and the lipid bilayer is denoted (two white lines). (b, c) Examples of 2D class averages (side-views) of spiked nanobicelles. (d) Montage of cryo-EM images of images (top-views) of spiked nanobicelles (I-IV). (e) Zoomed-in view of the spiked nanobicelle (panel I) to show details of HA top views. (f) Examples of 2D classes averages resulting from averaging individual HA spikes from the spiked nanobicelles displaying both side views (panels I, II) and top views (panels II, IV) of HA molecules. (g, h) Top and side views, respectively, of ab initio cryo-EM 3D reconstruction of HA spike. (i, j). Top and side views, respectively, of the docking of HA ectodomain trimeric coordinates into the 3D map. (k, l) Top-view (panel l) and side view (panel l) of the docking of H1 ectodomain coordinates (green ribbons) into a refined 3D map (light blue isosurface). Scale bar, 2nm. (m, n) Molecular models of spiked nanobicelles with membrane (gray) and constituent HA molecules with HA1 (red) HA2 (blue). Scale bars are denoted.

When observed in the top-view orientations, HA were arranged in rings (Fig. 2d, 2e). In some cases, top-views of HA molecular densities presented as 3-bladed propellers (Fig. 2e). 2D classification of individual spike images extracted from the SNBs revealed 2D classes with side views of peanut-shaped densities with a bottom-line density (Fig. 2f, panels I, II). Some other 2D classes presented densities as top-views of 3-bladed propellers (Fig. 2f, panels II, IV). Both an initial 3D map (ab initio) with coordinate docking (Fig. 2g-2j) and a 3D refined map with molecular docking of trimeric ectodomain coordinates (Fig. 2k, 2i) indicated that the constituent HA molecules were trimeric and in a prefusion conformation. Molecular models suggested that prefusion HAs are embedded in the lipid bilayer of the purified spiked nanobicelles (Fig. 2m, 2n).

### In vivo comparison between intramuscular immunization with Fluad or spike nanobicelles

To test if purified spiked nanobicelles (SNBs) were still immunogenic and protective mouse challenge studies were carried out to compare SNB (non-adjuvanted) and Fluad (MF59 adjuvanted) (Fig. 3a). SNBs were immunogenic producing significantly higher levels of antibodies to antigenically matched H1 HA compared to sera from mice that received saline (Fig. 3b, F(2,9)=666.6, p<0.0001). However, immunization with Fluad produced higher levels of H1 HA antibodies (p<0.0001) (Fig 3b). Immunization with SNBs or Fluad was able to provide 100% protection from an antigenically matched H1N1 challenge (Fig. 3c-3f). Although those immunized with SNB experienced approximately 10% weight loss during the challenge (day 3), mice recovered when compared to the saline group (Fig. 3d). Fluad immunized mice did not show body weight changes during challenge (Fig. 3f). When compared for differences in body weight on day 3 post challenge, Fluad did best in preventing weight loss (Fig. 3g, F(2,16)=52.62, p<0.0001), but SNB immunization attenuated the weight loss seen in the mice that received saline (p=0.0161) (Fig. 3g). When the experiment was repeated and mice were euthanized on the third day for tissue collection, mice that received saline had significantly more influenza virus in their lung tissue based on TCID50 titers than either of the vaccination groups (Fig. 3h, F(2,12)=4.897, p=0.0279, Fluad (p=0.0459) and SNB (p=0.0469).

**Fig. 3.**
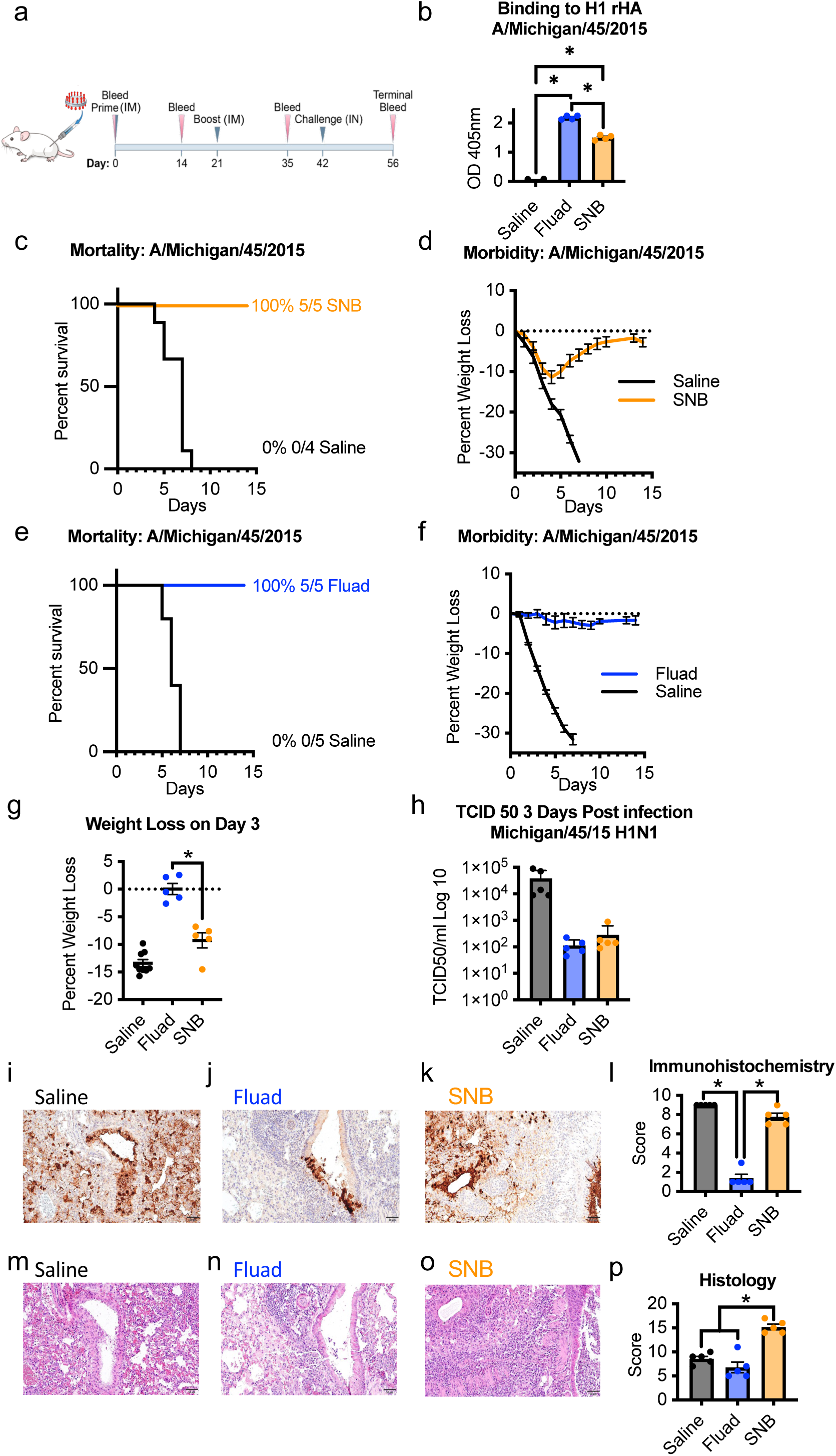
Protection efficacy of adjuvanted and non-adjuvanted spiked nanobicelles via intramuscular immunization against H1N1 influenza virus (2015) challenge. (a) Mice were intramuscularly immunized with adjuvanted commercial influenza vaccine Fluad and purified spiked nanobicelles (SNBs) (day 0 and 21) and intranasally challenged with A/Michigan/45/2015 H1N1 influenza virus at 10x the lethal dose (day 42). Mice were in immunization groups (N=5) consisting of Fluad, SNB, and saline. (b) Immunogenicity of Fluad (blue) and SNB (orange) assessed via ELISA to H1 HA 2015 with day 35 sera. (c, d) Survival curves and corresponding weight loss-curves for mice immunized with SNBs. (e, f) Survival curves and weight loss-curves for mice immunized with Fluad. (g) Comparison of weight-losses on day 3. (h) Comparison of mouse lung titers determined by 50% Tissue Culture infectious Dose (TCID50) for tissues taken day 3 post-H1N1 challenge with duplicate groups of mice. Graphs display mean with standard error of the mean bars. (i-k) Images show immunohistochemistry (IHC) against the influenza nucleoprotein protein for different vaccination groups and (l) scoring for images shown are from day 3 post H1N1 challenge for all groups. (m-o) Images of hematoxylin and eosin (H&E) staining of lung tissue sections for different vaccination groups and (p) histology scoring. Lung samples were collected from 2 mice per condition group and the experiment was replicated (N=4). All images are at 10x magnification and display 100 μm scale bars. Statistically significant differences are denoted with an asterisk to represent a significance level (p) less than 0.05.

Lung tissue samples were analyzed by immunohistochemistry and hematoxylin and eosin (H&E) staining to compare the level of viral antigen and pathology between SNB and Fluad (Fig. 3i-3p). The lung sections from saline treated mice had the highest level of viral antigen (i.e. brown stained areas) when compared to either the other treatment groups (Fig. 3i-3k) (Fig. 3l, F(2,12)=166.9, p<0.0001, Fluad (p<0.0001) and SNB (p=0.0487). The SNB vaccinated group had more viral staining than the Fluad vaccination group (p<0.0001), suggesting that following challenge there was increased viral replication in the lungs of SNB immunized mice compared to the lungs of Fluad immunized mice. However, by the third day the virus was no longer replicating differently between the groups based upon the TCID50 results (Fig. 3h). Another indication suggesting that SNB immunized mice had an increased viral load relative to Fluad immunized mice with subsequent clearance of infectious virus was the presence of inflammation (Fig. 3m-3p). Lung tissue slides showed significant differences in tissue damage scores between the treatment groups (Fig. 3p, F(2,12)=31.89, p<0.0001). Mice immunized with SNB had more inflammation than the saline (p=0.0002) and Fluad (p<0.0001) groups, while the Fluad and saline groups did not differ in their scores (p=0.2731). Thus, purified SNB elicited protective immunity, and the MF59 adjuvant appeared to aid in the reduction of both weight loss and lung pathology.

### In vivo comparison between intranasal immunization with spike nanobicelles and vaccines with different structural organizations of HA

While formulated with adjuvant, the Fluad vaccine is not recommended for intranasal immunization but the isolation of spiked nanobicelles (SNBs) allowed for different routes of administration to be examined. To determine if structural presentation of HA impacts intranasal immunogenicity, we selected influenza vaccines (SNB, Flublok, and Flumist) that differed in HA number and structural arrangements of HA complexes as shown by electron microscopy (Fig. 4a, 4b, 4c) (Supplementary Fig. 1, Supplementary Fig. 2). SNBs have rings of HA molecules arranged on a lipid bilayer disc (Fig. 4a). This differs from both the smaller starfish arrangements of HAs in Flublok (Fig. 6b) and the virion (i.e. virus particles) in Flumist (Fig 4c).

**Fig. 4.**
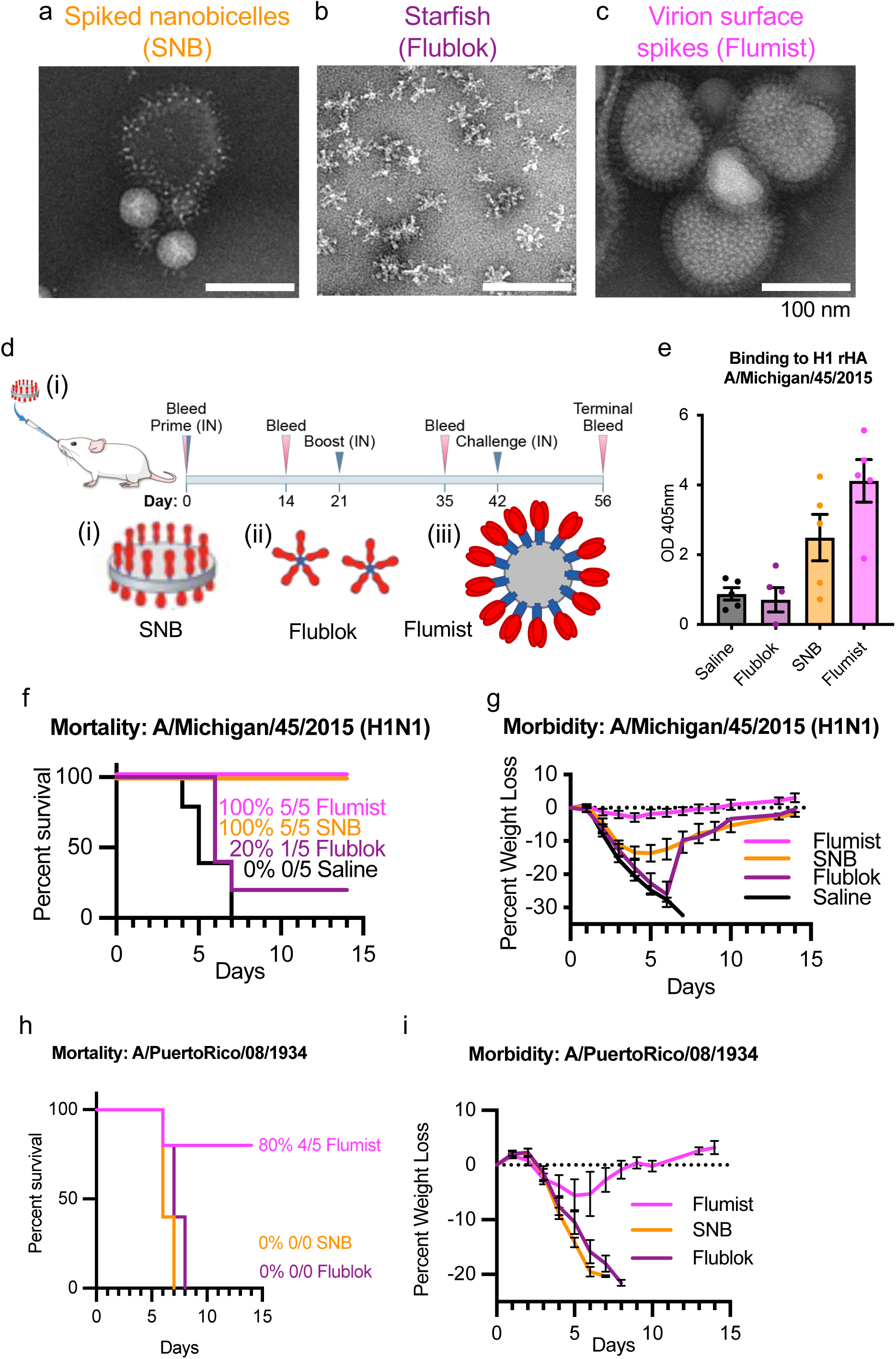
Comparison of protection efficacy of spike complexes organized as spiked nanobicelles, HA-starfish, and Virion (Surface spikes) via intranasal immunization against antigenically matched and mismatched H1N1 influenza virus challenge. (a, b, c) Negative-staining EM images of (a) spiked nanobicelles, (b) HA-starfish (Flublok) and (c) Virions (Flumist) of live-attenuated influenza vaccines (LAIV), respectively. (d) Intranasal immunization and challenge schedule for vaccines. Schematics (i, ii, iii) of each type of structural arrangement are shown with HA trimers with HA1 (red), HA2 (blue) and membrane gray. (e, f, g) Mice were intranasally immunized with purified spiked nanobicelles (SNBs) Flublok, Flumist, and Saline control (day 0 and 21) and intranasally challenged with A/Michigan/45/2015 H1N1 influenza virus at 10x the mouse lethal dose 50 (day 42). (e) Immunogenicity assessed via ELISA to H1 HA 2015 with day 35 sera. (f, g) Survival curves (panel f) and corresponding weight loss-curves (panel g) for the different groups of mice. (h, i) Survival curves (panel h) and corresponding weight loss-curves (panel i) for the different groups of mice challenged with A/Puerto Rico/08/1934 H1N1 influenza virus at 10x the mouse lethal dose 50. Immunization group colors are Saline (gray), Flublok (purple), Spiked nanobicelles (SNB) (orange) and Flumist (pink).

When groups of mice were primed intranasally and boosted intranasally with the different vaccines (Fig. 4d) there were differences in HA immunogenicity. Flumist, an FDA approved intranasal vaccine was a positive control for this route of administration and was immunogenic producing significantly higher levels of antibodies to antigenically matched H1 HA compared to sera from mice that received saline (Fig. 4e, F(3,16)=10.52, p=0.0005, Saline, p=0.0014) and those immunized with Flublok (p=0.0008). Flublok was not immunogenic via intranasal administration showing no difference from saline mice. While the SNB immunization group showed no difference from the saline group, they also did not differ from the Flumist immunization group (Fig. 4e), suggesting that SNB are eliciting an intermediary level of antibodies between the level of the negative control and the positive control.

Mice were then challenged with antigenically matched A/Michigan/45/2015 (H1N1), both Flumist and SNB provided 100% protection from viral challenge. In contrast, Flublok only provided 20% protection (Fig. 4f). Immunization with Flumist reduced weight loss (Fig. 4g). Flumist had significantly less weight loss at three days after challenge than all other groups (F(3,16)=21.11, p<0.0001) (Supplementary Fig. 10a) and the lowest virus titers in lung tissues (Supplementary Fig. 10b). However, the high variability of titers between individual animals in the groups limited the detection of differences that were statistically significant (Supplementary Fig. 10b). The Flumist group had less viral antigen staining in the lungs at day 3 post infection when compared to all other groups (Supplementary Fig. 10c-10g) but there were no group differences in inflammation (Supplementary Fig. 10h-10l). This suggest that at day 3 the SNB immunized mice have not yet begun to recover from the viral challenge, as there was no sign of lowered replication rate (TCID50) or viral clearance (inflammation) suggesting that the time course for protection differs between intranasal and intramuscular vaccination.

### Testing for protection against antigenically mismatched H1N1 via intranasal vaccination

To test for pan-H1 protection, mice were intranasally immunized with SNB, Flublok, and Flumist and then challenged with antigenically mismatched A/Puerto Rico/8/1934 (H1N1). All vaccines failed to provide 100% protection (Fig. 4h, 4i). There was variation in the level of protection with Flumist at 80%, SNB at 0%, and Flublok at 0% (Fig. 4h). Flumist immunized mice challenged with antigenically mismatched A/Puerto Rico/8/1934 (H1N1) lost approximately 5% of their body weight (Fig. 4i) in contrast to mice challenged with antigenically matched A/Michigan/45/2015 (H1N1) which did not change from baseline (Fig. 4g).

### Antigenic variation of H1 Has

To further understand antigenic variation between the H1 HA 2015-like vaccine antigens and the 1934 challenge virus, the H1 HA sequences were compared along with a 2009 pandemic virus (Supplementary Fig. 11). The majority of the HA mutations observed between the 2015 and 2009 H1 strains occur in the HA1 regions (16 mutations) versus HA2 regions (3 HA2 mutations) (Fig. 5a, red residues). Sequence identity was 98% (Supplementary Fig. 11). When the 2015 HA strain was compared to a classical seasonal influenza, A/Puerto Rico/8/1934, changes in amino acids were present in both HA1 (81 mutations) and HA2 (15 mutations) (Fig. 5b). Sequence identity was 81% (Supplementary Fig. 11). Thus, bioinformatics analysis indicated that these H1 HAs could be used as models of H1 HA antigenic drift because the 2015, 2009, 1934 H1 HA proteins displayed antigenic drift/variation.

**Fig. 5.**
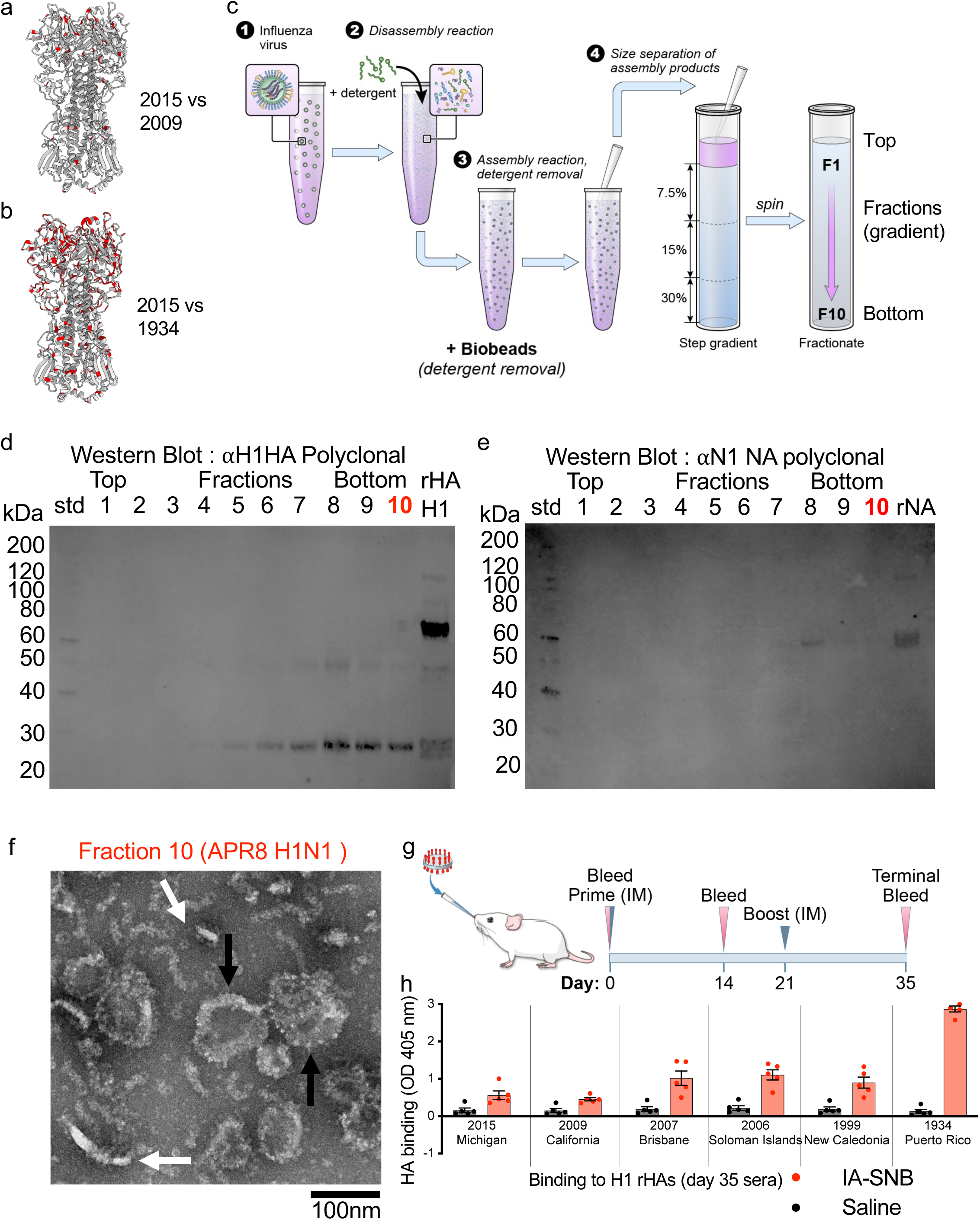
In vitro assembly of 1934 H1 spiked nanobicelles (SNBs) and testing the elicitation of pan H1 immunogenicity via intranasal administration. (a, b) H1 HA trimeric ectodomain structure of H1 HA 2015 (PDB 7KNA) with residue differences from H1 HA (2009) (panel a) and residue differences from H1 HA (1934) (panel b) indicated in red. (c) Schematic of the process to produce in vitro assembled spiked nanobicelles (IA-SNBs). (Step 1) Influenza virus (A/Puerto/Rico/8/24 H1N1) was disassembled into structural components via (Step 2) detergent-mediated lysis. (Step 3) Detergent was removed via the addition of biobeads. (Step 4) Detergent removal resulted in the assembly products (e.g. spiked nanobicelles) that were purified via gradient centrifugation. Steps are denoted by numbers. (d) Probing for influenza viral H1 HA protein in gradient fractions by western blot analysis. Recombinant H1 HA (rHA HA) was a positive control. (e) Probing for influenza viral neuraminidase (NA) protein in gradient fractions by western blot analysis. Recombinant neuraminidase (rNA) was a positive control. Fraction 10 is denoted by bold underlining because it was subjected to further analysis by electron microscopy. Fraction 10 should contain large (megadalton) complexes based on its gradient density. (f) Negative-staining electron microscopy of fraction 10 from sucrose gradient displaying in vitro assembled spike nanobicelles (IA-SNB). Each hemagglutinin (HA) appears as a spike with multiple spikes emanating from a central disc of lipid. Scale bar, 100 nm. (g) Intranasal immunization schedule for invitro assembled spiked nanobicelles (IA-SNBs) of 1934 H1 HA. (h) ELISA used to probe for pan-H1 binding antibodies in day 35 sera. Saline is a control with H1 HA recombinant proteins from influenza viruses from the following years: 2015, 2009, 2007, 2006, 1999, 1934, corresponding to A/Michigan/45/2015 (H1N1), A/California/07/09 (H1N1), A/Brisbane/59/2007 (H1N1), A/Solomon Islands/3/2006 (H1N1), A/New Caledonia/20/1999 (H1N1) and A/Puerto Rico/08/1934 (H1N1) influenza viruses.

### In vitro assembly of spiked nanobicelles from H1 1934 influenza virus

An in vitro assembled spiked nanobicelles (IA-SNB) protocol was developed (Fig. 5c), to begin virus was dissembled (i.e. disrupted) with detergent, followed by subsequent removal of detergent with biobeads and purification of IA-SNB by gradient centrifugation. After centrifugation, fractions were taken from the top to the bottom of the gradient (fractions 1 to 10) (Fig. 5c). Western blot analysis indicated that fraction 10 (based on density of the gradient) should have megadalton sized structures and had HA protein (Fig. 5d). Influenza NA was undetectable in fraction 10 (Fig. 5e). Negative-staining electron microscopy confirmed the presence of IA-SNB in fraction 10 (Fig. 5f). In side-views IA-SNB appeared as flat discs displaying HA spikes (Fig. 5f, white arrows). In some top-views IA-SNB appeared as rings of protruding spikes (Fig. 5f, black arrows). Subsequent lipidomics analysis confirmed enrichment of long chain ceramides and hexosylceramides in the IA-SNBs compared to the lower density fractions, similar to Fluad-SNBs (Supplementary Fig. 5b-5c). Thus, based upon the western blot, negative stain EM, and lipidomics, this protocol was successful in producing HA in vitro assembled spiked nanobicelles (IA-SNBs) comparable to Fluad derived spiked nanobicelles (SNBs).

### Breadth of immunogenicity for intranasal administration of in vitro assembled spiked nanobicelles

Using HA protein probes, we found that intranasally administered 1934 H1 IA-SNB elicited pan-H1 HA binding antibodies to antigenically divergent H1 HAs (Fig. 5g, 5h). Day 35 sera showed binding via ELISA to a panel of antigenically drifted H1 HA proteins spanning from 2015 to 1934 (Fig. 5h). The highest binding was for the antigenically matched H1 1934. There was a trend of decreased binding going forward in years to 2015 for antigenically mismatched H1 HAs (Fig. 5h). However, binding activities to 2015 and 2009 H1 HAs were above the saline control (Fig. 5h).

### Intranasal administration of in vitro assembled spiked nanobicelles provide pan-H1 protection but not intramuscular administration

A mouse influenza virus challenge model was used to determine if 1934 H1 IA-SNB immunizations could provide pan-H1 protection via intramuscular (Fig. 6a-6g) and intranasal immunization (Fig. 6h-6n). Following intramuscular immunization, the 1934 IA-SNBs provided 100% protection (survival) against the antigenically matched 1934 H1N1 virus (Fig. 6b) with no decrease in body weight (Fig. 6c). For antigenically mismatched viruses the protection was 40% for 2009 H1N1 (Fig. 6d, 6e) and 80% for 2015 H1N1 (Fig. 6f, 6g) influenza challenges viruses, respectively. Even those mice that survived 2009 and 2015 challenge had increased weight loss (morbidity) (Fig. 6d-6g).

**Fig. 6.**
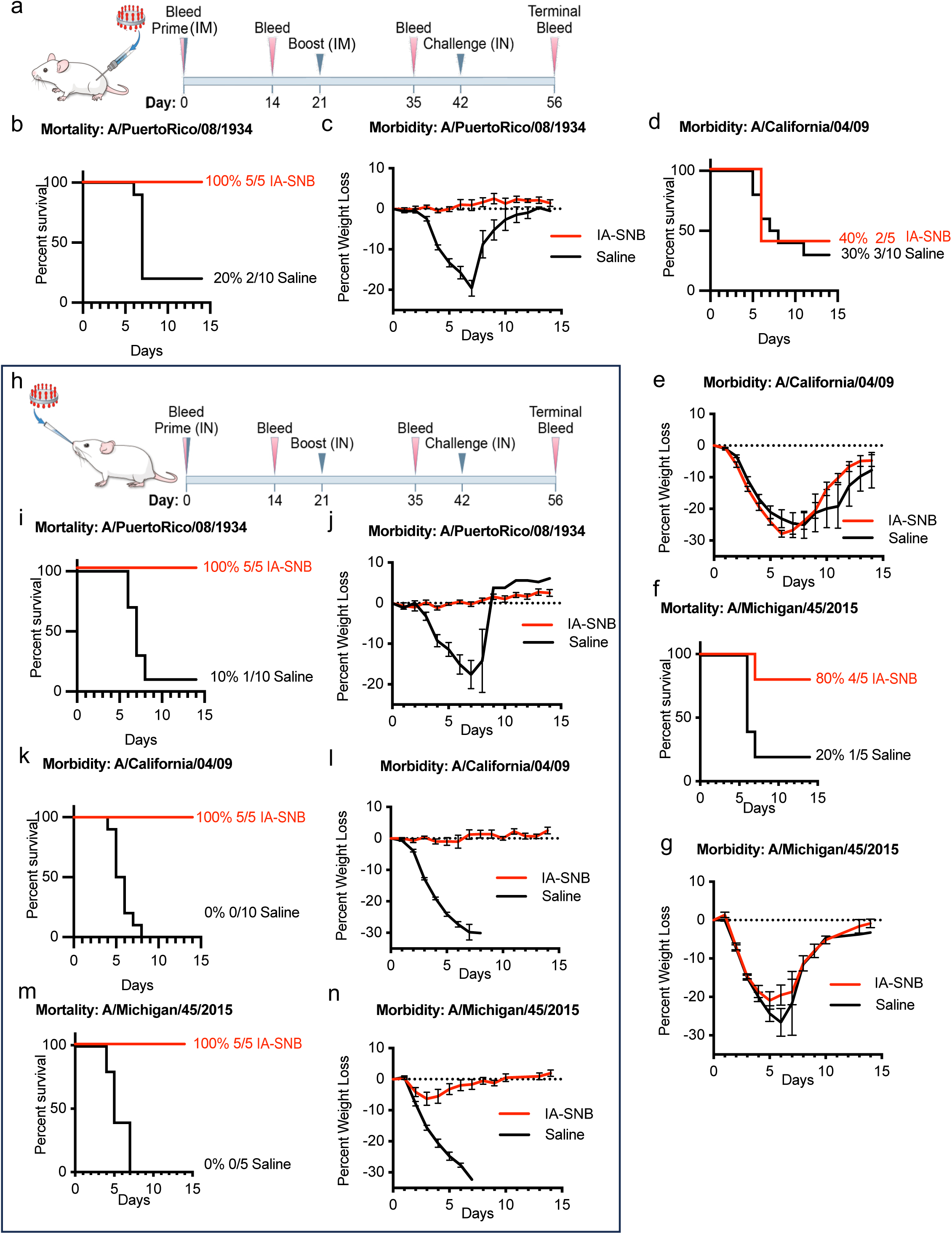
Pan H1 protection efficacy of in vitro assembled spiked nanobicelles via intramuscular and intranasal immunization against antigenically drifted H1N1 influenza viruses. (a) Intramuscular immunization and lethal challenge schedule for in vitro assembled spiked nanobicelles (IA-SNBs). Mice were intramuscularly immunized (day 0 and 21) with IA-SNBs and saline control. IA-SNBs contained H1 HA 1934-like antigens. Groups of mice were intranasally challenged on day 42 with antigenically matched A/Puerto Rico/08/1934 H1N1 or antigenically mismatched A/California/07/09 H1N1 or /Michigan/45/2015 H1N1 influenza viruses. (b) Survival curves and (c) weight loss-curves for immunized mice challenged with A/Puerto Rico/08/1934 H1N1. (d) Survival curves and (e) weight loss-curves for immunized mice challenged with A/California/07/09 H1N1. (f) Survival curves and (g) weight loss-curves for immunized mice challenged with A/Michigan/45/2015 H1N1. (h) Intranasal immunization and lethal challenge schedule for in vitro assembled spiked nanobicelles (IA-SNBs). Survival and corresponding weight loss-curves are shown for the following influenza viruses: (i, j) A/Puerto Rico/08/1934 H1N1, (k, l) A/California/07/09 H1N1, and (m, n) A/Michigan/45/2015 H1N1.

In sharp contrast, intranasal administration of the 1934 IA-SNBs provided 100% protection and reduction in weight loss for both antigenically matched and mismatched H1N1 influenza virus challenges (Fig. 6h-6n). For the antigenically matched challenge, the intranasal 1934 IA-SNB provided 100% protection and no weight loss against lethal challenge with A/Puerto Rico/8/1934 H1N1 (Fig. 6i, 6j).

When challenged with the antigenically mismatched pandemic virus, A/California/04/2009 H1N1, mice survived at 100% with no weight loss (Fig. 6k, 6l). Although minor weight loss was observed for the antigenically mismatched challenge with A/Michigan/45/2015 H1N1, intranasal administration of the 1934 H1 IA-SNB provided 100% protection (Fig. 6m, 6n).

The protection provided by IA-SNB immunizations appeared to be antibody mediated, because naïve mice were partially protected (80%) from viral challenge after passive-transfer of antibodies from mice immunized with IA-SNB (Supplementary Fig. 12). Antibody profiles were produced for antibody class and IgG subclasses in sera. Intranasal IA-SNB immunization elicited higher levels of IgG2a and IgG3 antibodies when compared to intramuscular IA-SNB immunizations (Supplementary Fig. 13a, 13b).

Flumist vaccination elicited the highest level of IgA and IgG3, but intranasal IA-SNB immunization also elected higher levers of IgG2a and IgG3 than those elicited by immunization with SNB purified from Fluad (Supplementary Fig. 13c).

## Discussion

Influenza HA is the primary target for elicitation of neutralizing and protective antibodies against influenza virus, but it is prone to antigen drift ^39,40^. Elicitation of antibodies targeting more conserved areas of HA have shown broad efficacy against various antigenically variable influenza viruses ^41^.

However, production of cross-reactive anti-HA antibodies appear limited in response to seasonal influenza viruses requiring yearly updating to vaccines. Some current efforts to elicit immune responses to conserved HA epitopes focus on the use of a recombinant scaffold of symmetric nanoparticles to display ectodomains in the forms of trimeric HA ectodomains ^20,29,42^, engineered stabilized trimer HA stems ^21,22,30^ or HA1 heads ^19^. The current hypothesis is that nanoparticle spacing of antigens at 5-10 nm can increase the activation of B-cells^43,44^. Nanoparticle display increases the multivalency of antigen and these particles have been shown to improve the immune response to the displayed epitope ^20,29,42^ ^45^. A limitation of previous structure-guided nanoparticle studies of HA ectodomain regions is that they usually rely on designing and screening HA constructs that fold and display epitopes correctly on non-membrane containing nanoparticles. These nanoparticle platforms have also required the addition of an adjuvant to improve immune responses and have been administered via intramuscular immunization ^20–22,29,30,42–44^. Furthermore, these particles are often designed with symmetrical presentation of HA, such as octahedral^20–22^ or icosahedral^42^ symmetry, which is necessary for the inserted epitopes to be displayed correctly on the nanoparticle.

Our structure-guided approach to synthesize spiked nanobicelles (SNB) suggest that there may be other nanoplatform options for eliciting broadly reactive and protective antibodies to antigenically variable viruses, such as influenza. We found that even though SNB do not have symmetrical insertions of HA like a ferritin nanoparticle, they still display a high number of HA on a single SNB, an average of up to 60 molecules of HA trimeric proteins (Fig. 1, Fig. 2, Fig. 4a). And while SNB do not have strict symmetry, they are arranged in an ordered array with 5-10 nm spacing between HA molecules (Fig. 2a-2e), the spacing reported for B-cell activation ^44,46^. These constituent HA molecules appear to be in in a prefusion state (Fig. 2f-2i), which is needed for the elicitation of broadly protective antibodies whose epitopes are formed only on the trimeric prefusion state of HA ^21,22,30,42,47^.

When further characterize by lipidomic analysis, an increase in very long chain sphingolipids was associated with the SNB vaccine preparation in both Fluad SNB (Fig.1h) and A/PR8 derived IA-SNB (Supplementary Fig. 5c). Sphingolipids are involved in forming specialized ordered membrane microdomains called lipid rafts. These rafts play crucial roles in cell signaling, protein trafficking, and membrane organization^48^. In addition, sphingolipids are involved in multiple aspects of immunity, including immune cell activation, migration, and inflammatory responses^49^. Interestingly, lipid rafts serve as assembly and budding sites for influenza viruses and hemagglutinin concentrates in lipid raft microdomains ^50^. This suggests that the source of sphingolipids in SNB are structurally ordered plasma membrane regions participating in viral budding that get included as part of the viral membrane. The depletion of sphingolipids in the lower density upper fractions that contain the MF59 adjuvant suggest that these sphingolipids are not derived from the MF59 adjuvant (Fig. 1h).

Our ability to reproduce sphingolipid-enriched SNB structures from virus in the absences of MF59 adjuvant (in vitro-assembled spiked nanobicelles (IA-SNB)) further strengthens the proposition that the sphingolipid components originate in the viral membrane and are not associated with the adjuvant. The recruitment of these specific lipids into influenza virions and the subsequent inclusion of these lipids in the HA-rich SNB structures suggests that this sphingolipid-rich lipid composition is important for assembling full-length HA molecules. Sphingolipids and lipids with longer saturated acyl-chains form more rigid ordered structures, and these lipids might be a key driver of the formation of the stable disc bilayer of the spiked nanobicelles (Fig. 2a, Fig. 5F, Supplementary Fig. 7). In future experiments it will be important to control sphingolipid content with regard to both headgroup and acyl-chain length with the IA-SNB production protocol to determine if changes in sphingolipid formulation result in alterations to the structural display of HA. As we’ve hypothesized that the viral membrane is the source of the sphingolipid and needed for synthesis of the SNB and IA-SNB structure, it will be interesting to test the assembly protocol on other subtypes like influenza H3N2 and influenza B HAs to see if they will also produce lipid discs.

In addition to different viral subtypes, the method of viral growth could affect the structure of the IA-SNB. Traditionally virus used in vaccine manufacturing has been egg-propagated despite the possibility of mutations in HA that can arise from egg-adaption of influenza viruses^51^. Non-egg cell-based methods have emerged for the production of commercial influenza vaccines ^5,15,52,53^. However, egg-based production still remains a major means for the production of commercial influenza vaccines, like Fluad (Supplementary Fig. 1). Additionally, our current production of IA-SNB utilized an egg-propagated virus as our starting material from which we obtained HA. Immunization with either production method of lipid discs, by intramuscular or intranasal routes, provided protection against an antigenically matched challenge (Fig. 3c, 4f, 6b, 6i), suggesting that this particular structural organization of HA might abrogate the effect of any egg adapted mutations.

If this structural platform can mitigate egg mutations, our IA-SNB protocol could have broader applications to vaccine development because egg-grown influenza virus particles themselves, without their associated glycoproteins, are being used as display platforms and delivery vehicles. For example, reverse genetics has been used to display chimeric HA on influenza virus particles^54^. In this work, a chimeric HA had an H1 stalk and an H5 globular head domain (cH5/1 HA) displayed on influenza virus that was grown in 10-day-old specific-pathogen-free embryonated chicken eggs^54^. In other influenza egg work, influenza was engineered to express and display the spike protein of SARS-CoV-2. Newcastle disease virus (NDV) has been engineered to express that SARS spike protein. NDV vaccines were purified from eggs and used to develop both inactivated and live virus vaccines against SARS-CoV-2^55,56^. It would be interesting to apply the SNB in vitro assembly assay to such NDV viruses to determine if spiked nanobicelles (SNB) of other viral glycoprotein spikes can be produced. This would indicate a general method for producing viral glycoprotein nanostructures that could be used as both intramuscular and intranasal vaccines (Fig. 6). Our assembly assay could be scaled up using the egg-based production of the influenza vaccine manufacturers.

When challenged with an antigenically mis-matched challenge virus only the intranasal immunization with IA-SNB resulted in pan H1 protection (Fig. 5, Fig. 6). The inability of the SNB from Fluad to produce pan H1 protection with the same administration paradigm (Fig. 4h, 4i) could be due to differences in the HA displayed on the surface of the lipid disc. HA on the SNB was isolated from Fluad synthesized from inactivated virus while the IA-SNB originated from disrupted but not inactivated virus. Chemical inactivation of HA could alter the amino acid residues forming epitopes of HA. This could lead to diminished potency of the vaccine, lower hemagglutination titers and loss of immunogenicity of key antigenic epitopes^57^. This means that inactivated virus may not induce as broad immune response as desired. The HA on these platforms also differ on when the H1 virus was circulating in human populations, Fluad contains a modern H1 from after the 2009 pandemic while the IA-SNB utilized an older H1 from before the pandemic (1934). While we showed the location of mutations in HA between the viruses (Fig. 5a-5b and Supplementary Fig. 11), it is possible that there is directionality where some mutations are more difficult to overcome moving backwards in time, leading to the differences we see in survival.

Another possible explanation for the lack of pan H1 protection from immunization with SNB from Fluad could be the presence of multiple HA types in the starting material. Fluad being a commercially available influenza vaccine, it is required to include HA from H1, H3 and B viruses (Yamagata lineage and Victoria lineage), while our IA-SNB began with only H1 HA. Since Fluad contains 4 HA antigens quantification of the H1 HA in our SNB was difficult and overall protein level was used to determine concentration, this calculation means that the IA-SNB could have been administered at a higher H1 HA concentration than the SNB from Fluad. Future experiments with SNB should include a range of protein concentrations of immunizations to determine if administering more protein could produce pan H1 protection.

There are also differences in the amount of SNB and IA-SNB protein administered based upon the route chosen, intranasal or intramuscular. Mice immunized via intramuscular injections received four times the amount of lipid disc as their intranasal counterparts. This may be one reason that the intranasal vaccinations resulted in more variability in immune responses than the intramuscular route (Fig. 3b and Fig. 4e). Alternatively, the variability could be due to differences in retention of the vaccine at the site, the bleeds being conducted systemically giving an inaccurate representation of mucosal antibodies, or a shift in antibody isotype elicited by intranasal vaccination as the detection for the assay was specific for the IgG isotype.

If a shift in isotype is occurring with intranasal administration of SNB, there would be an increase in the levels of IgA detected in the sera of mice. However, these levels were less than that for the intranasal vaccine Flumist (Supplementary Fig. 13). These results suggest that circulating IgA levels in the sera might not be a good surrogate for lung IgA or that the live-attenuated virus in Flumist is better in eliciting IgA. Nevertheless, SNB and IA-SNB vaccinations can produce circulating antibodies that convey protection to naïve mice against antigenically matched H1N1 viruses (Fig. 3c, Fig. 6b). Sera from intranasal immunization with the IA-SNB had increases in the levels of both IgG2a and IgG3a when compared to intramuscular immunization (Supplementary Fig. 13).

IgG2 antibodies, like IgG2a can enhance the complement activity of sera to aid in control of influenza infection^58^. IgG3 is also effective at activating the complement system. IgG3 antibodies can recognize and neutralize antigenically drifted influenza viruses more effectively than other IgG subtypes with increased bivalent binding to influenza HAs and SARS-CoV-2 spikes^59^. One speculation is that the arrangement of the neighboring HAs on the spiked nanobicelles might aid in the promotion of bivalent binding antibodies. Further studies on how alterations to the HA structural arrangement and the size and composition of spiked nanobicelles (SNB) can affect the antibody class and subclass distribution following immunization are needed. It will be important to understand how different antibody classes and subclasses correlate with eliciting broadly protective immune responses to antigenically variable influenza viruses.

In conclusion, although influenza vaccines in this study contained full-length HA proteins, there were differences in the number of HA molecules per particle (multivalency), spacing, and epitope display within HA complexes. Vaccines differed in their ability to elicit pan H1 immunogenicity and protection via the intranasal route of administration. Our results suggest that vaccine formulations that specifically structure HA antigen into spiked nanobicelles (SNB) can offer improved multivalency and elicitation of cross-reactive antibodies with increased breath of protection. HA-based SNB formulated as novel intranasal HA subunit vaccines could aid in the development of more efficacious influenza vaccines for seasonal antigenic drift, pandemic antigenic shift and potentially universal influenza vaccine development. More broadly, our structure-guided approach to the synthesis of in vitro assembled spiked nanobicelles (IA-SNB) can be applied to other viral glycoproteins to produce intranasal vaccine platforms for other respiratory viruses such as RSV and SARS coronaviruses.

## Methods

### Fluad adjuvant CryoEM

Quantifoil R2/1 300 mesh grids (Ted Pella) were glow discharged for 25 sec and 15 mA using a PELCO easiGlow, then samples were plunge frozen with a Leica EM GP set to 80% humidity at 25 °C. Three μl sample was applied to the grid, blotted for 1.2 sec, then plunge frozen in liquid ethane. Frozen grids were loaded on a Gatan model 626 cryo holder and inserted into a Tecnai 12 electron microscope operative at 100 kV (Thermofisher). SerialEM ^60^ software controlled a Gatan OneView camera to acquire low-dose images, for a total dose per image of approximately 80 e-/Å2. And 3.9 Å /pixel.

### Negative stain EM

Formvar/carbon substrate on 300 mesh copper grids (Ted Pella) were glow discharged for 25 sec and 15 mA using a PELCO easiGlow, then 3 μl sample was applied. After 30 sec, sample was blotted from the grid, then the grid was immediately subjected to 3 cycles of dH2O rinsing and blotting, then applying 1% uranyl acetate negative stain for 1 min. The grid was blotted obliquely until dry. After air-drying for 30 min or longer, a Tecnai 12 electron microscope and OneView camera were used to acquire images. Low-dose mode was not used as the negative stain was stable on the timescale needed for image acquisition.

### Single Particle CryoEM

C-flat R 2/2 200 mesh grids were glow discharged for 10 sec with O2/H2 mix (80%/20%) using a Gatan Solarus, then samples were plunge frozen with a Leica EM GP2 set to 95% humidity at 5 °C. Three μl sample was applied to the grid, blotted for 3 sec, then plunge frozen in liquid ethane. Frozen grids were loaded on a Thermofisher Titan Krios using a 20-eV energy filter slit width and a Gatan K3 camera operating in super-resolution mode and with correlated-double sampling mode enabled. EPU software was used to acquire 5,217 movies, at 54 e-/Å2 total dose and 0.66 Å/pixel. Images were processed in CryoSparc ^61^ computing on either NIAID HPC Skyline, or the AWS cloud.

For reconstruction of individual HA proteins, patch-based motion correction was used to align and sum movies into images, while binning 4x. The contrast transfer function (CTF) for each micrograph was estimated using CTFFIND4. Manual picking initially obtained 2,889 particles using a box size of 216 Å, consisting of both top and side views of HA on nanobicelles. 2D classification was used to select the subset of 1,812 particles consistent with HA structure, then Cryosparc was used to generate an ab-initio model. Homogenous Refinement was used to refine the 3D model with C3 symmetry imposed.

For 2D class averaging of intact nanobicelles, 454 side views were selected from 300 micrographs using a box size of 686 Å. Cryosparc 2D classification was performed with 5 classes, of which 4 were well resolved. For top views, 71 nanobicelles were separately picked from 1300 micrographs using a box size of 1,240 Å. Spiked nanobicelles with larger diameters tended to prefer top view orientations and not side views. Two rounds of 2D classification resulted in 4 classes that were well resolved.

### Cryo-EM reconstruction

The cryo-EM data was processed using Cryosparc v 4.5.3. Visualization, analysis, and segmentation for mask generation were performed using ChimeraX v 1.6.1. 5,217 movies were aligned, and contrast transfer functions (CTF) were calculated for the resulting micrographs. 4,067 micrographs (78%) with CTF-fit better than 5Å and an ice-thickness range of 0.9 – 1.1. Manually picked nanobicelle-bound HA particles, were used to generate templates and 4.8 million initial particles were template picked. After exclusion of 2D classes containing ice or crowded particles 902,682 particles with a 320-pixel box were aligned within a mask of the highest-resolution regions of the HA protein.

### Sequence alignment and image rendering

Amino acid sequences were pairwise aligned with NCBI BLAST ^62^. Protein structure renderings and rigid-body fitting of atomic coordinates to EM density were produced with UCSF Chimera ^63^. Nanobicelle renderings were created using Blender 3D (www.blender.org). Scale bars and distance measurements were made with FIJI software ^64^. PDB files used in this study were downloaded from the Protein Data Bank: https://www.rcsb.org. These included the following PDBs: 7kna, 4fnk, and 2rfu.

### Isolation of Spiked nanobicelles from Fluad

Spiked nanobicelles (SNB) were isolated from Fluad vaccine by sucrose gradient centrifugation. Fluad (500μl) was layered on top of a 7.5%, 15%, and 30% sucrose step gradient. The sample was spun for 6 hrs at 8°C in a SW 55 Ti rotor at 32,500 rpm (100,000 × g) within a XL-70 Ultracentrifuge (Beckman Coulter, Fullerton, CA). Fractions (500 μl) were collected from the top to bottom of the gradient. SDS-PAGE gels and Western blots were used to analyze the protein composition of the gradient fractions and to identify fractions that contained HA. Electron microscopy was used to identify HA fractions that contained adjuvant particles and spiked nanobicelles.

### In vitro assembly of spiked nanobicelles from virus

In vitro assembled spiked nanobicelles (IA-SNB) were assembled from detergent disrupted influenza virus:A/Puerto Rico/8/34 (H1N1) (Charles River) that was subsequently treated with detergent removing Bio-Beads SM-2 resin (Bio-Rad). The virus disassembly reaction consisted of influenza virus (1 ml)(2mg/ml) that was mixed with 50 μl of Triton X-100 (10 % stock). The reaction was incubated at 30 °C for 1 hr. 500μl of the disassembly reaction was incubated with 0.5g of prewetted and degassed Bio-Beads for 24 hours at 4°C. The solution was then separated from Bio-Beads using a micropipette. 500 μl of the assembly reaction was then placed on a sucrose step gradient and centrifugation was as detailed above for Fluad. SDS-PAGE gels and Western blots were used to analyze the protein composition of the gradient fractions and to identify fractions that contained HA. Electron microscopy was used to identify HA fractions that contained in vitro assembled spiked nanobicelles. SNBs were buffered exchanged into PBS via dialysis cassettes (ThermoFisher Scientific) and concentrated by centrifugal filtration (EMD Millipore).

### Metabolite and Lipid Sample Preparation

For all liquid chromatography mass spectrometry (LCMS) methods LCMS grade solvents were used. All samples were immersed in 0.4 mL of ice-cold methanol. To each sample 0.4 mL of water and 0.4 mL of chloroform were added. Samples were agitated under refrigeration for 30 minutes and centrifuged at 16k xg for 20 min to establish biphasic layering. 400 µL each of the bottom layer (organic) was collected. The organic layer was dried down under vacuum and resuspended in an equivalent volume of 5 µg/mL butylated hydroxytoluene in 6:1 isopropanol:methanol for targeted bulk lipidomics.

### Liquid Chromatography Mass Spectrometry

LCMS grade water, acetonitrile, and all buffer additives were purchased through Fisher Scientific. Bulk lipidomics was performed as previously described ^65^ with a shortened gradient and polarity switching. A LD40 X3 UHPLC (Shimadzu Co.) and a 7500 QTrap mass spectrometer (AB Sciex Pte. Ltd.) were used for separation and detection. Lipids were separated by lipid class on a Water XBridge Amide column (3.5 μm, 3 mm X 100 mm) with a 9-minute binary gradient from 100% 5 mM ammonium acetate, 5% water in acetonitrile apparent pH 8.4 to 95% 5 mM ammonium acetate, 50% water in acetonitrile apparent pH 8.0. All lipids were detected using scheduled MRMs that leveraged fatty acid product ions, headgroup product ions, or neutral loss ion combinations that were conserved within each lipid class.

### Mice immunizations

Immunogenicity and challenge studies were performed under mouse protocols approved by the Animal Care and Use Committee (ACUC) at the National Institute of Allergy and Infectious Diseases (NIAID). Experiments were conducted exclusively in female mice based upon their reported higher immunogenic response following vaccination ^66–68^. Initial immunogenicity experiments used 10 BALB/c mice (Taconic Biosciences) aged 8-10 weeks that were randomly assigned to either SNB (5) or saline (5). Intramuscular immunizations (IM, 50μL/leg) occurred on day 0 (prime) and day 21 (boost) at a concentration of 0.68mg/ml (SNB, Fluad) or 0.25mg/ml (IA-SNB, 1934 H1). Mice were weighed weekly to ensure health with bleeds occurring on Day 0, Day 14, and Day 35 to track immunogenicity.

Mice used in protection studies were immunized at a similar schedule as the immunogenicity experiment with the route of administration being either IM or intranasal (IN, 25ul). On day 42 mice were anesthetized and intranasal challenged with 10xMLD50 (50% Mouse Lethal Dose) of influenza virus, A/Michigan/45/2015, A/California/04/2009 ^69^ or A/Puerto Rico/45/1934.

Survival experiments used five mice per group. Mice were weighed daily and observed twice daily for survival criteria (animals were euthanized if they lost ≥30% of their initial body weight for A/Michigan/45/2015, A/California/04/09 and ≥20% for A/Puerto Rico/45/1934) until Day 56, when all surviving mice were euthanized. Differences in survival rates were compared using a Kaplan-Meier survival analysis and comparison of survival curves by Log-rank (Mantel-Cox) test. Replicate mouse studies were conducted for organ collection on Day 45, 3 days post-challenge. Experimental groups remained the same as the protection studies with the addition of a sham inoculation group that received intranasal administration of saline supplemented with 0.1% bovine serum albumin (BSSBSA) on Day 42 instead of virus challenge.

### Immunoassays (ELISA)

Enzyme linked immunosorbent assay (ELISA) was conducted to determine the immunogenicity of the vaccine candidates. Sera samples collected on Day 35 were tested against recombinant H1 HA antigen. Between all steps plates were washed with PBS plus 0.1% Tween 20 (405 TS ELISA Plate Washer, Agilent Technologies). Antigen was applied to 96-well plates and incubated overnight at 4°C (1.25 mg/mL). The next day, plates were blocked (1% Omniblok, AmericanBio, Inc., and 0.1% Tween20 in PBS) for 30 minutes at room temperature before mouse serum samples were plated at a 1:125 dilution. Plates were then incubated at 37°C with a HRP conjugated secondary antibody (goat anti-mouse IgG (H+L), Thermo Fisher Scientific). Colorimetric detection occurred at room temperature for 15 minutes and measured for absorbance at 405 nm (1-Step ABTS, Thermo Fisher Scientific). Analysis of variance (ANOVA) tests were conducted and Tukey’s multiple comparisons test to determine group differences were conducted for those with a significant main effect, reported as the F statistic with degrees of freedom within and between.

### Immunoassays

For immunoblots (western blots), samples were separated by SDS-PAGE before transfer to nitrocellulose membranes via the iblot transfer system (Thermo Fisher Scientific). Denatured samples loaded onto the SDS-PAGE were fractions taken from the sucrose gradient. Blots were probed with primary antibody to HA (Protein Sciences), NA (GeneTex), M1 (HB64), or NP (HB65). Matching fluorescent-labeled secondary antibodies, goat anti-rabbit IgG and goat anti-mouse IgG (Thermo Fisher Scientific), were applied via lateral flow (ibind system, Thermo Fisher Scientific). Membranes were visualized on the c600 imager (Azure biosystems).

### Tissue culture infectious dose (TCID50)

96-well plates were seeded with 3 × 10⁴ MDCK cells per well. The next day, in 50μl of DMEM with TPCK-Trypsin (0.5 ml/ml) the virus was serially diluted twofold and added to the cells in eight replicates, with two negative controls included cell control with DMEM without TPCK-Trypsin, and the second with DMEM with TPCK-Trypsin. After incubation at 37 °C for 1h, 50μl more of DMEM with FBS and antibiotic was added. Then, the infected cells were incubated at 37 °C for 72 h. After this time, the media was removed and cells were washed with PBS and fixed with methanol, stained with Crystal Violet. Further, the plates were visualized using a Cytation 7 reader and Gen5 software (Agilent Technologies Inc.). The analysis was performed by normalizing the confluence percentage of the cell control, DMEM with TPCK, to 100% for each row and TCID50/ml was quantified based on the Spearman and Karber method using a calculator developed by Marco Binder, Heidelberg University. Histology and immunohistochemistry were carried out as previously detailed ^23,70^.

## Acknowledgements

This work was supported by the Intramural Research Program of the National Institute of Allergy and Infectious Diseases. We also thank Vinod Nair, Cindi Schwartz, Elizabeth Fischer, and Rick Huang for help in cryo-EM data collection. Joseph Marcotrigiano provided access to Cloud Computing with Amazon Web Services (AWS). This work utilized the computational resources of the NIH HPC Biowulf cluster (http://hpc.nih.gov), and the Office of Cyber Infrastructure and Computational Biology (OCICB) High Performance Computing (HPC) cluster at the National Institute of Allergy and Infectious Diseases (NIAID), Bethesda, MD.

## Author Contributions Statement

MM designed the in vivo experimental studies, ELISA experiments and western blots. JG and AD designed the EM experiments. NK, WP, JG, AD and AH collected and analyzed EM data. SM-P grew viral reagents and conducted TCID50 experiments. DW and AH optimized the sucrose purification protocol. KB and DA conducted and interpreted the pathology experiments. EB and BS conducted and interpreted the lipidomic experiments. MM, JG, and AH designed the study and experiments and wrote the paper. All authors contributed to editing the paper.

---

**Supplementary Fig. 1. Commercial influenza vaccines used in this study.** Details are provided for the 3 commercial vaccines studied: Fluad, Flublok, and Flumist. (a) The total HA concentration and the concentration of HA components such as H1 or H3 HA subtypes are denoted, as well as (b) production and inactivation methods.

**Supplementary Fig. 2. Negative-staining electron microscopy of commercial influenza vaccines used for intranasal challenge studies.** (a) Fluad spiked nanobicelles (white arrows). A large adjuvant particle is denoted by a black arrow. (b) A field of Fluad spiked nanobicelles. (c) Flublok HA-starfish and (d) Flumist virions of live-attenuated influenza vaccines (LAIV). Scale bars, 100 nm. Contrast is with protein density as white.

**Supplementary Fig. 3. Sucrose gradient separation of MF59 adjuvant and purification of spiked nanobicelles from Fluad.** (a) Milky Fluad layered on top of the gradient pre-centrifugation. (b) Post-centrifugation with a milky ban at the top of the gradient. The top and bottom of the gradient and fraction #1 and fraction #10 are denoted.

**Supplementary Fig. 4. Western blot analysis of fractions 1 to 10 of the sucrose gradient used for purification of spiked nanobicelles from Fluad.** (a) Western blot of fractions taken from the top to the bottom of the gradient to probe for the present of HA protein via anti-H1 HA polyclonal antibody. (b, c, d) Similar analysis as in panel a, but with individual blot and antibodies for NA, M1 and NP as follows: (b) anti-NA polyclonal antibody, (c) anti-M1 polyclonal antibody, (d) anti-NP polyclonal antibody. Fraction 10 (orange) contained the spiked nanobicelles and is denoted. Controls were recombinant NA, M1, and NP proteins and lysozyme. Molecular weight standards (kDa) are denoted.

**Supplementary Fig. 5. Lipidomic analysis from sucrose gradients.** (a) Analysis of Fluad fraction 10 (i.e. spiked nanobicelles) enrichment for lipid components. (b) Analysis of A/PR8 fraction 10 (i.e. in vitro assembled spiked nanobicelles) enrichment for lipid components. (c) Lipid profile with lipid classes that increase (positive values) and decrease (negative values) for fraction 10 (in vitro assembled spiked nanobicelles) when compared to fraction 3.

**Supplementary Fig. 6. Cryo-electron microscopy of purified spiked nanobicelles purified from Fluad.** (a) Observation of spiked nanobicelles complexes (white arrows). Select HAs on the complexes are denoted (white circles). Clusters of individual HA molecules are denoted (white brackets). The framed area is shower at larger scale in panel b. (b) Zoomed in view of area in panel a with spiked nanobicelles (white arrow) and isolated HA molecules (white brackets). Scale bars are indicated: 100nm, 10nm. Contrast is with protein density as black.

**Supplementary Fig. 7. 1D profile analysis of spiked nanobicelles from cryo-electron microscopy.** (a) An example of a 2D-class average of a spiked nanobicelle (Fluad) with protruding spikes and a lipid bilayer. The yellow line represents the line profile used to calculate a trace. Scale bar, 10 nm. (b) 1D density profile trace with the distance between lipid bilayer peaks (4.3 nm) and thickness of the lipid bilayer (10.6 nm) denoted. Contrast is with protein density as white.

**Supplementary Fig. 8. 2D-class averages of top-views of spiked nanobicelles from cryo-electron microscopy.** (a-d). Montage of individual 2D class averages. Scale bar, 10 nm. Contrast is with protein density as white.

**Supplementary Fig. 9. Comparison of the docking of different HA coordinates into 3D maps of HA.** (a, b) Top and side views, respectively, of ab initio cryo-EM 3D reconstruction of HA spike. (c, d). Top and side views, respectively, of the docking of H1 HA ectodomain trimeric coordinates (PDB 7kna) into the 3D map (gray). (e, f). Top and side views, respectively, of the docking of H3 HA ectodomain trimeric coordinates (PDB 4fnk) into the 3D map (gray). (g, h). Top and side views, respectively, of the docking of influenza B HA ectodomain trimeric coordinates (PDB 2rfu) into the 3D map (gray). The 3D map is shown as an isosurface rendering (gray) and coordinates are ribbon diagrams with H1 (red), H3 (blue) and influenza B HA (yellow). Scale bar, 10nm. (i, j) Top-view (panel i) and side view (panel j) of the docking of H1 ectodomain coordinates (green ribbons) into a refined 3D map (light blue isosurface). Scale bar, 2nm. (k) Corresponding FSC curves (estimated resolution, 4.1 Angstroms).

**Supplementary Fig. 10. Pathology comparisons of vaccines via intranasal immunization against H1N1 influenza virus (2015) challenge.** (a) Comparison of weight-losses on day 3. Graphs display mean with standard error of the mean bars. (b) Comparison of mouse lung titers determined by 50% Tissue Culture Infectious Dose (TCID50) for tissues taken day 3 post-H1N1 challenge with duplicate groups of mice. (c-f) Images show immunohistochemistry (IHC) against the influenza nucleoprotein protein for different vaccination groups and (g) scoring of images shown are from day 3 post H1N1 challenge for all groups. (h-k) Images of hematoxylin and eosin (H&E) staining of lung tissue sections for different vaccination groups and (l) histology scoring. Lung samples were collected from 2 mice per condition group and the experiment was replicated (N=4). All images at 10x magnification and display 100 μm scale bars. Immunization group colors are Saline (gray), Flublok (purple), Spiked nanobicelles (SNB) (orange) and Flumist (pink). Statistically significant differences are denoted with an asterisk to represent a significance level (p) less than 0.05.

**Supplementary Fig. 11. Sequence comparison of H1 HA proteins from 1934, 2009, and 2015 influenza viruses.** (a) GenBank accession numbers for the H1 HA protein sequences. (b) Sequence alignment of H1 HA sequences. Sequence regions corresponding the HA1 (red) and HA2 (blue) are denoted with amino acids differences colored (black). Arginine (R) is underlined marking the separation of HA1 and HA2 regions. (c) comparison of the number of mutations and sequence identity between A/Puerto Rico/8/1934 and A/Michigan/15/2015 H1N1 influenza viruses.

**Supplementary Fig. 12. Passive-transfer of vaccine sera of in vitro assembled spiked nanobicelles (IA-SNB) for protection from H1N1 challenge**. (a) Survival curves for mice that were intranasally challenged with influenza virus (A/Puerto Rico/8/1934 (H1N1)) 24 hours after intraperitoneally (IP) transfer of 400ul of saline sera (black line) or sera collected from mice immunized with 1934 H1 HA spiked nanobicelles (IA-SNB) (red line). (b) The corresponding weight-lost curves for panel a, with IA-SNB vaccine sera (red line) and saline negative-control (black line).

**Supplementary Fig. 13. Analysis of relative amount of different antibody classes and subclasses in different vaccine sera a day 35.** (a) Sera from mice intramuscularly immunized with spiked nanobicelles from Fluad (SNB)(orange) or in vitro assembled spiked nanobicelles (IA-SNB) (red) of 1934 H1 HA. (b) Sera from mice intranasally immunized with spiked nanobicelles from Fluad (SNB)(orange) and in vitro assembled spiked nanobicelles (IA-SNB) (red). (c) Comparing antibody classes and subclasses of sera from intranasally administered Flumist (pink) to intranasally administered Fluad SNB (orange) and IA-SNB (red).

