## Supplementary Figures for "Structure-guided assembly of an influenza spike nanobicelle vaccine provides pan H1 intranasal protection"

**a** Vaccine Components

| Name | Lot # | Total HA | HA/Strain | H1N1 Strain | H3N2 Strain | Influenza B-Victoria | Influenza B-Yamagata |
|---|---|---|---|---|---|---|---|
| Fluad | 346374<br>346360 | 60ug/0.5ml | 15ug/0.5ml | A/Victoria/2570/2019<br>IVR-215 | A/Darwin/6/2021<br>IVR-227 | B/Austria/1359417/2021<br>BVR-26 | B/Phuket/3073/2013<br>BVR-1B |
| Flublok | UJ909AA | 180ug/0.5ml | 45ug/0.5ml | A/Wisconsin/588/2019 | A/Darwin/6/2021 | B/Austria/1359417/2021 | B/Phuket/3073/2013 |
| Flumist | PM2440 | | $10^{6.5\text{-}7.5}$ FFU/0.2ml | A/Victoria/1/2020 | A/Norway/16606/2021 | B/Austria/1359417/2021 | B/Phuket/3073/2013 |

**b** Vaccine Manufacturing

| Name | National Drug Code | Manufacturer | Vaccine Type | Valency | Inactivation | Method of Disruption | Reported Purification Method |
|---|---|---|---|---|---|---|---|
| Fluad | 70461-122-04 | Seqirus | Subunit, Adjuvanted | Quadrivalent | Formaldehyde | Cetyltrimethylammonium bromide | Zonal centrifugation |
| Flublok | 49281-722-88 | Protein Sciences | Recombinant | Quadrivalent | | Triton X-100 | Column chromatography |
| Flumist | 66019-309-10 | MedImmune | Attenuated virus | Quadrivalent | | | Ultracentriifugation |

**Supplementary Fig. 1. Commercial influenza vaccines used in this study.** Details are provided for the 3 commercial vaccines studied: Fluad, Flublok, and Flumist. (a) The total HA concentration and the concentration of HA components such as H1 or H3 HA subtypes are denoted, as well as (b) production and inactivation methods.

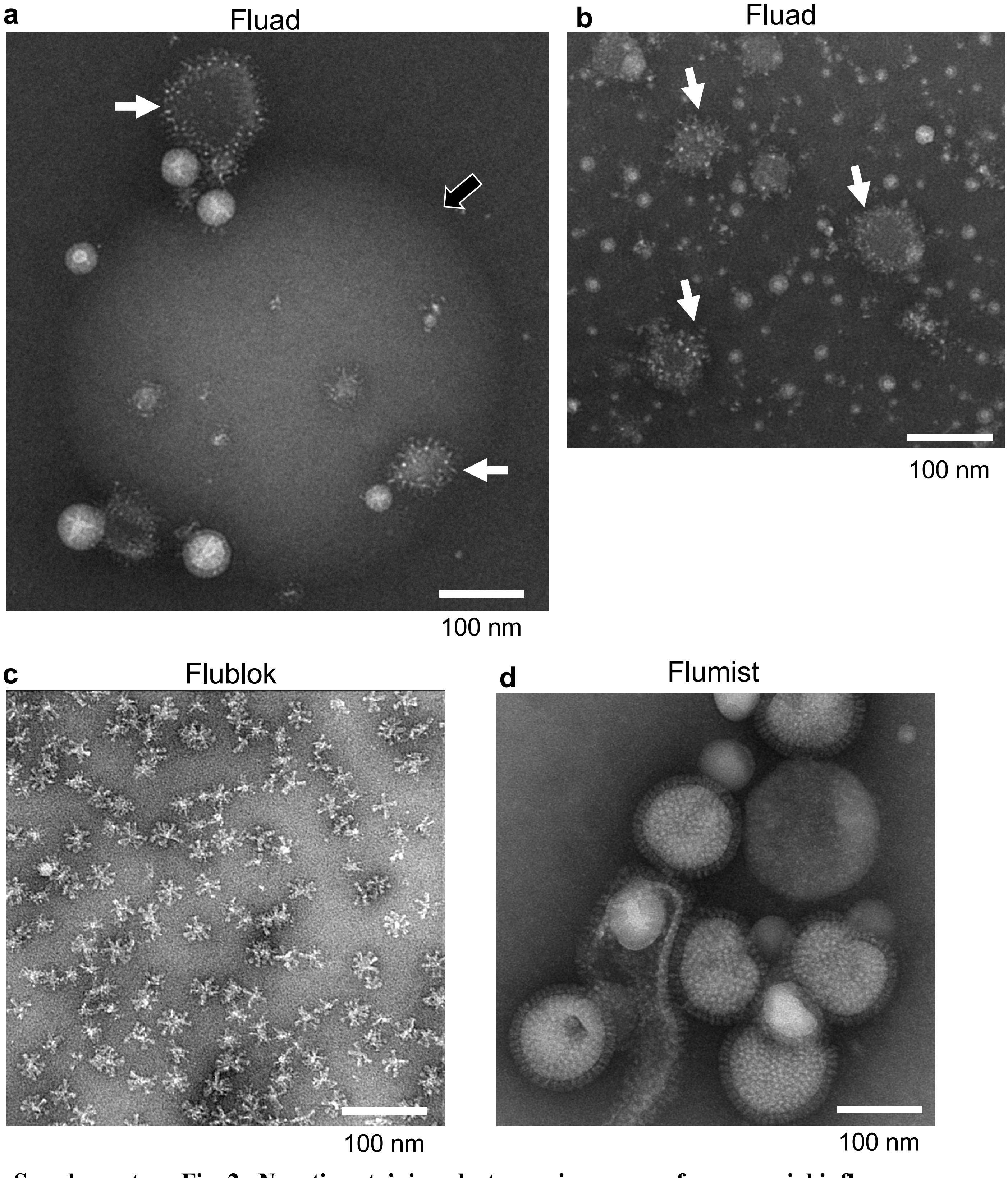


**Supplementary Fig. 2. Negative-staining electron microscopy of commercial influenza vaccines used for intranasal challenge studies.** (a) Fluad spiked nanobicelles (white arrows). A large adjuvant particle is denoted by a black arrow. (b) A field of Fluad spiked nanobicelles. (c) Flublok HA-starfish and (d) Flumist virions of live-attenuated influenza vaccines (LAIV). Scale bars, 100 nm. Contrast is with protein density as white.

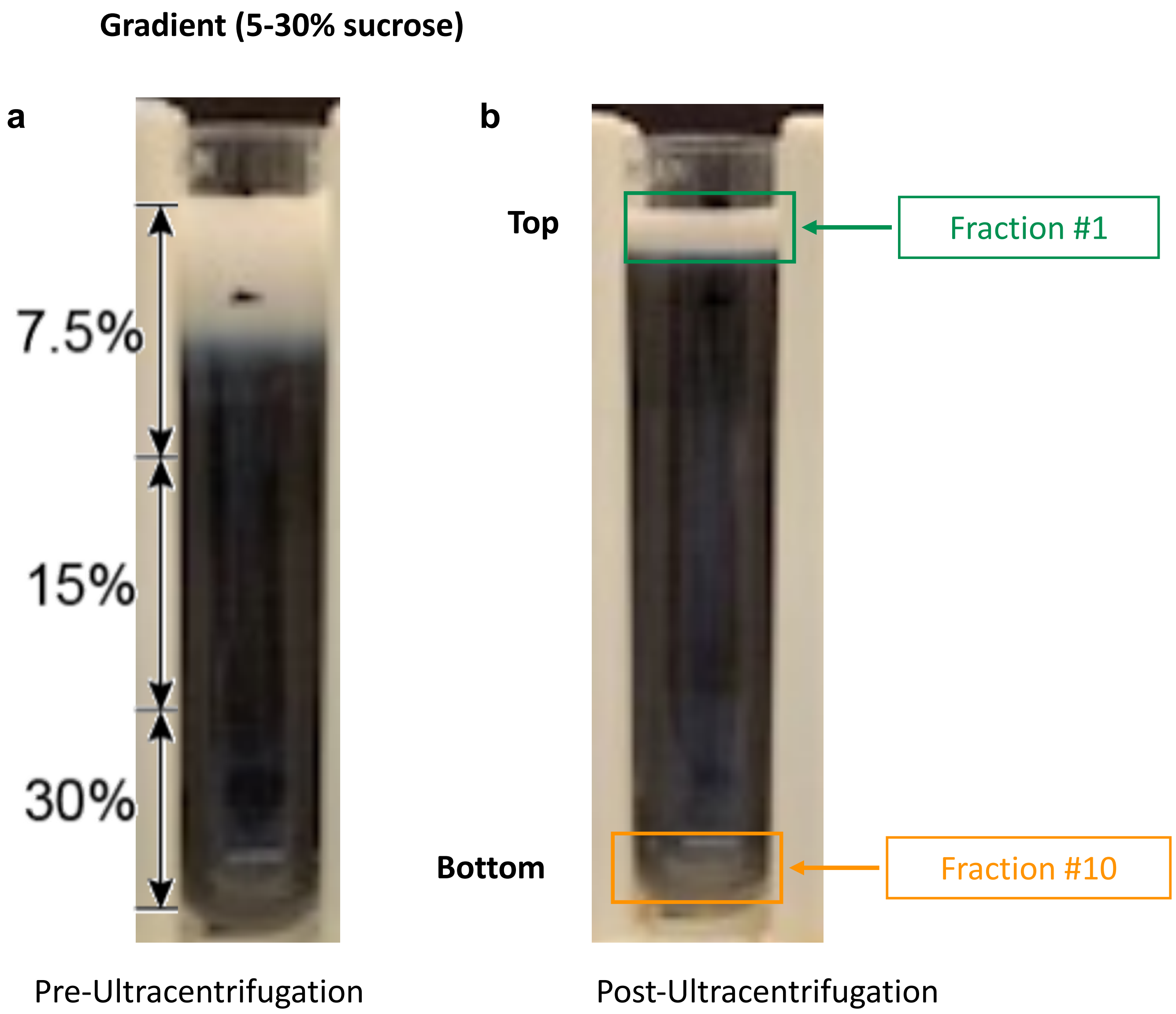


**Supplementary Fig. 3. Sucrose gradient separation of MF-59 adjuvant and purification of spiked nanobicelles from Fluad.** (a) Milky Fluad layered on top of the gradient pre-centrifugation. (b) Post-centrifugation with a milky ban at the top of the gradient. The top of the bottom of the gradient and fraction #1 and fraction #10 are denoted.

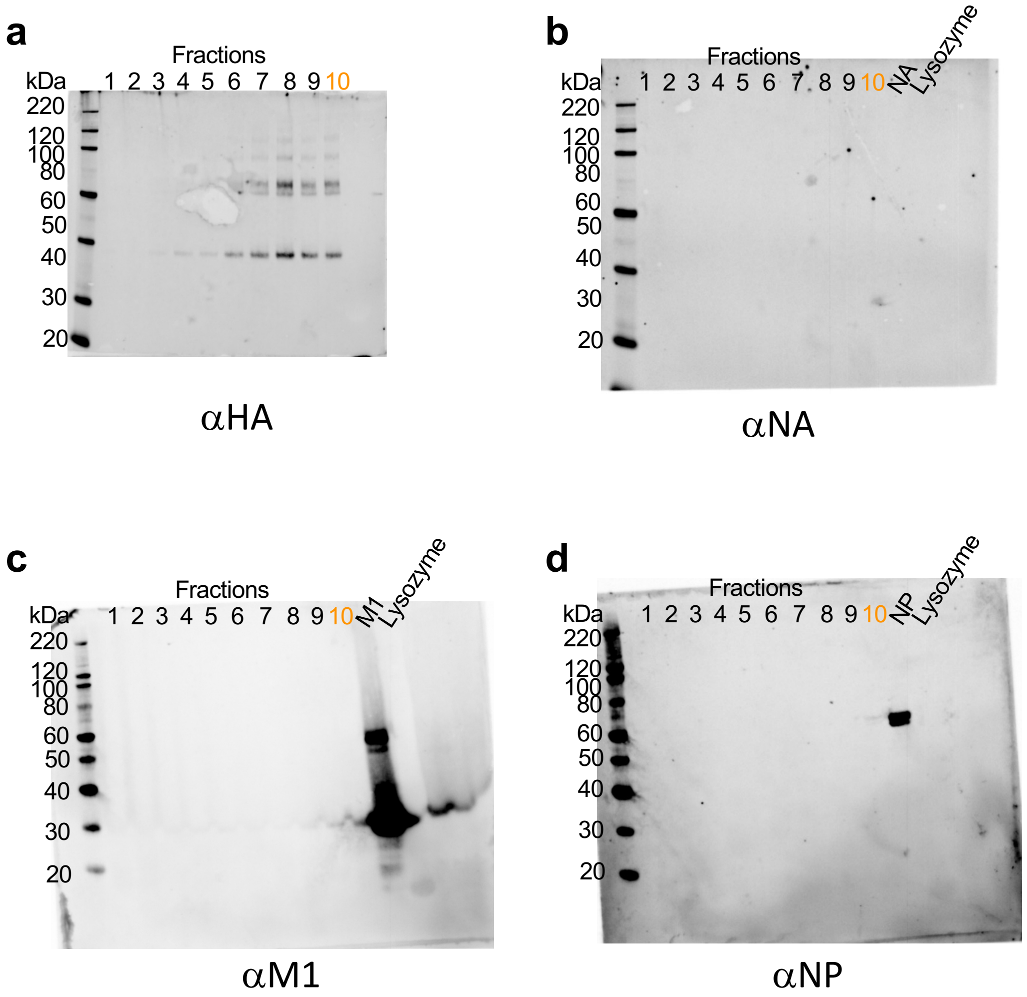


**Supplementary Fig. 4. Western blot analysis of fractions 1 to 10 of the sucrose gradient used for purification of spiked nanobicelles from Fluad.** (a) Western blot of fractions taken from the top to the bottom of the gradient to probe for the present of HA protein via anti-H1 HA polyclonal antibody. (b, c, d) Similar analysis as in panel a, but with individual blot and antibodies for NA, M1 and NP as follows: (b) anti-NA polyclonal antibody, (c) anti-M1 polyclonal antibody, (d) anti-NP polyclonal antibody. Fraction 10 (orange) contained the spiked nanobicelles and is denoted. Controls were recombinant NA, M1, and NP proteins and lysozyme. Molecular weight standards (kDa) are denoted.

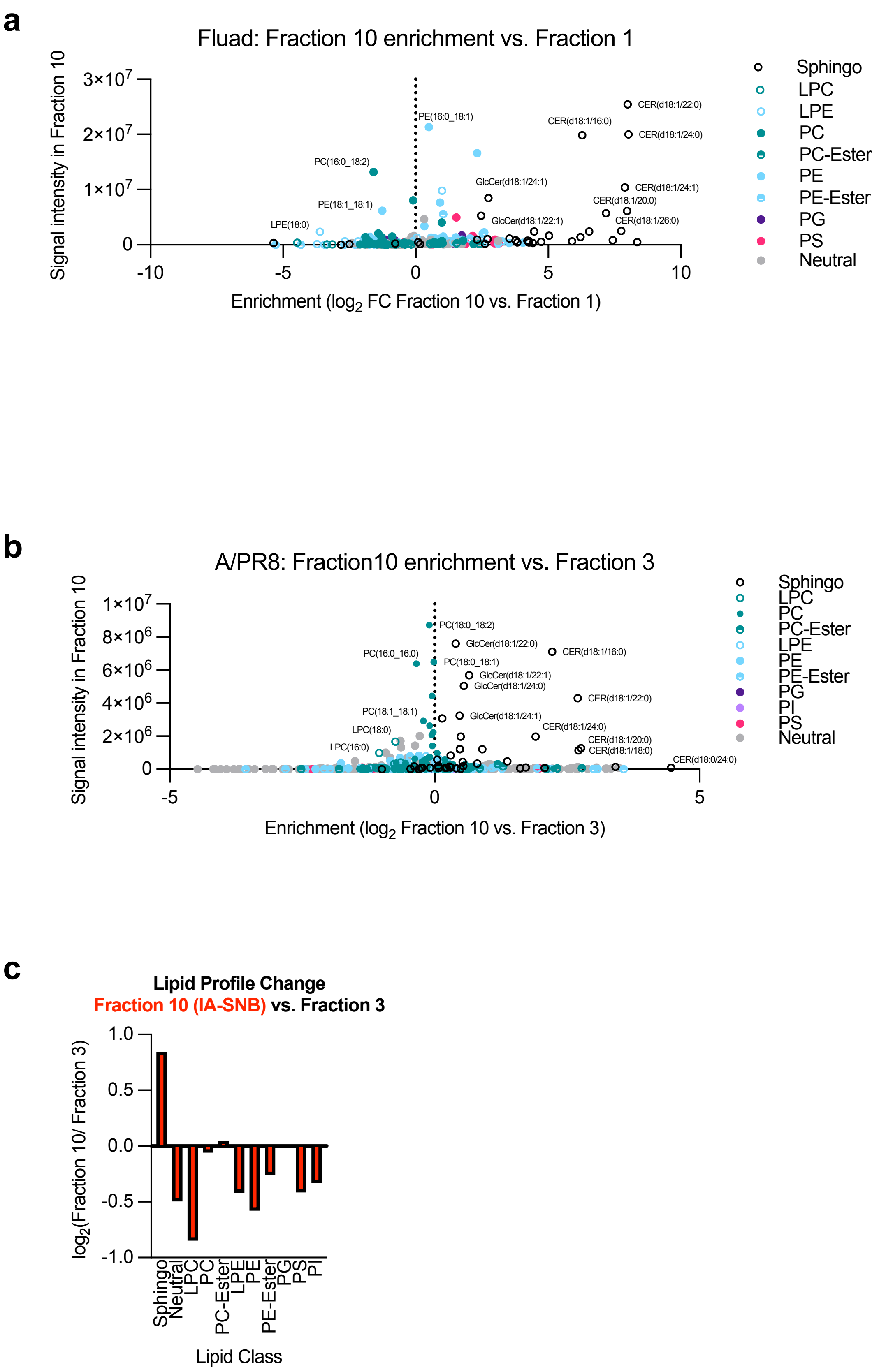


**Supplementary Fig. 5. Lipidomic analysis from sucrose gradients.** (a) Analysis of Fluad fraction 10 (i.e. spiked nanobicelles) enrichment for lipid components. (b) Analysis of A/PR8 fraction 10 (i.e. in vitro assembled spiked nanobicelles) enrichment for lipid components. (c) Lipid profile with lipid classes that increase (positive values) and decrease (negative values) for fraction 10 (in vitro assembled spiked nanobicelles) when compared to fraction 3.

**a**

**b**

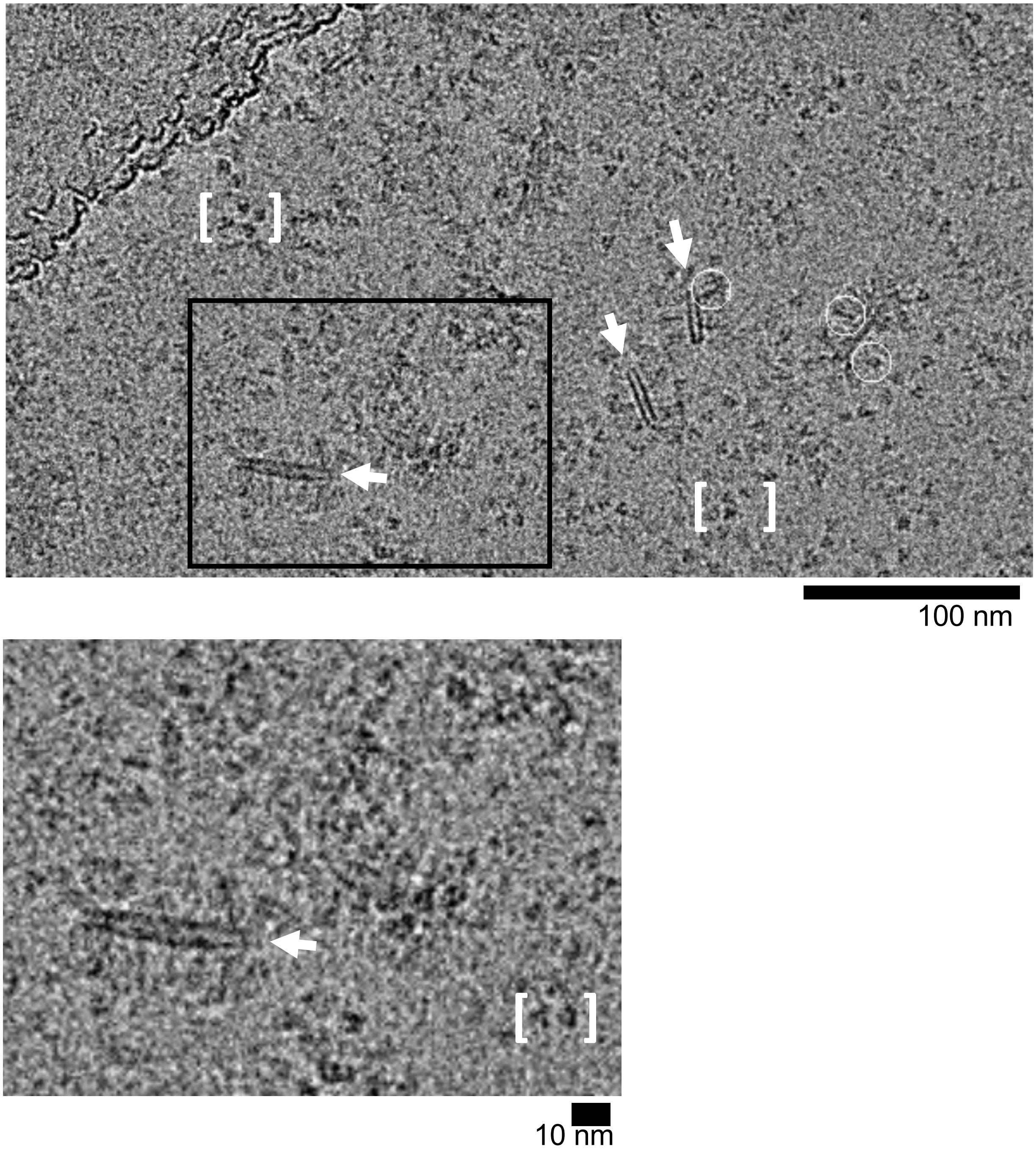


**Supplementary Fig. 6. Cryo-electron microscopy of purified spiked nanobicelles purified from Fluad.** (a) Observation of spiked nanobicelles complexes (white arrows). Select HAs on the complexes are denoted (white circles). Clusters of individual HA molecules are denoted (white brackets). The framed area is shower at larger scale in panel b. (b) Zoomed in view of area in panel a with spiked nanobicelles (white arrow) and isolated HA molecules (white brackets). Scale bars are indicated: 100nm, 10nm. Contrast is with protein density as black.

a

b

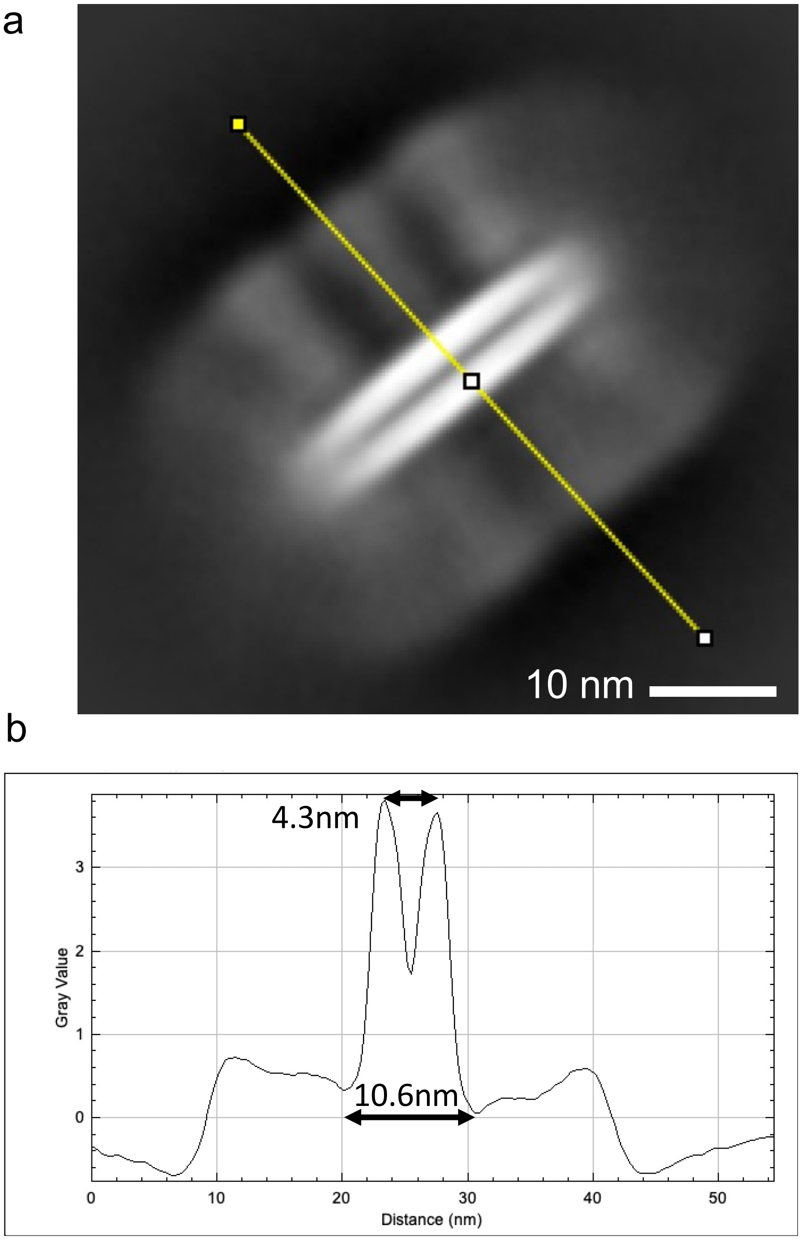


**Supplementary Fig. 7. 1D profile analysis of spiked nanobicelles from cryo-electron microscopy.** (a) An example of a 2D-class average of a spiked nanobicelle (Fluad) with protruding spikes and a lipid bilayer. The yellow line represents the line profile used to calculate a trace. Scale bar, 10 nm. (b) 1D density profile trace with the distance between lipid bilayer peaks (4.3 nm) and thickness of the lipid bilayer (10.6 nm) denoted. Contrast is with protein density as white.

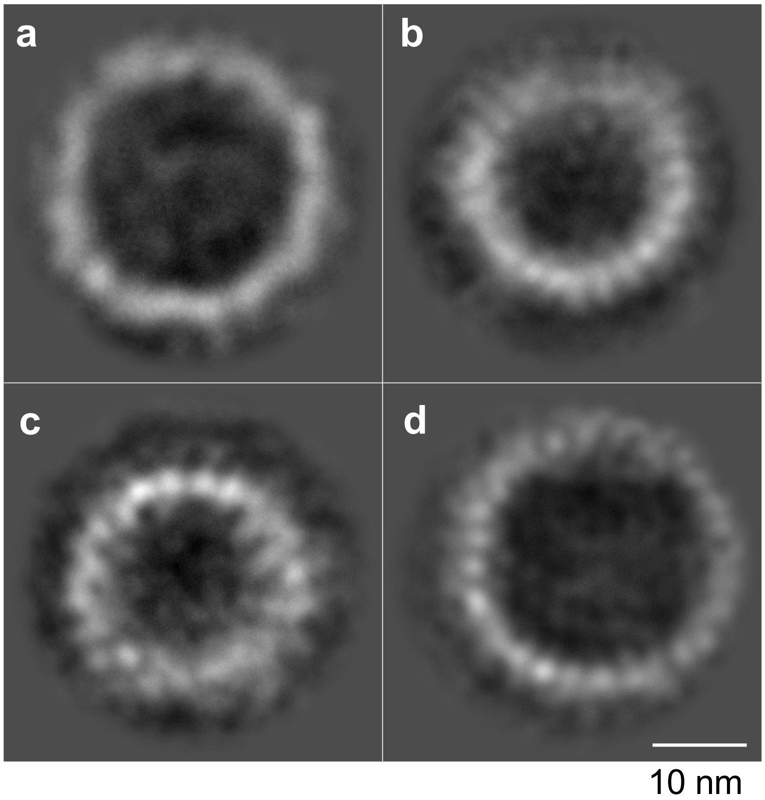


**Supplementary Fig. 8. 2D-class averages of top-views of spiked nanobicelles from cryo-electron microscopy.** (a-d). Montage of individual 2D class averages. Scale bar, 10 nm. Contrast is with protein density as white.

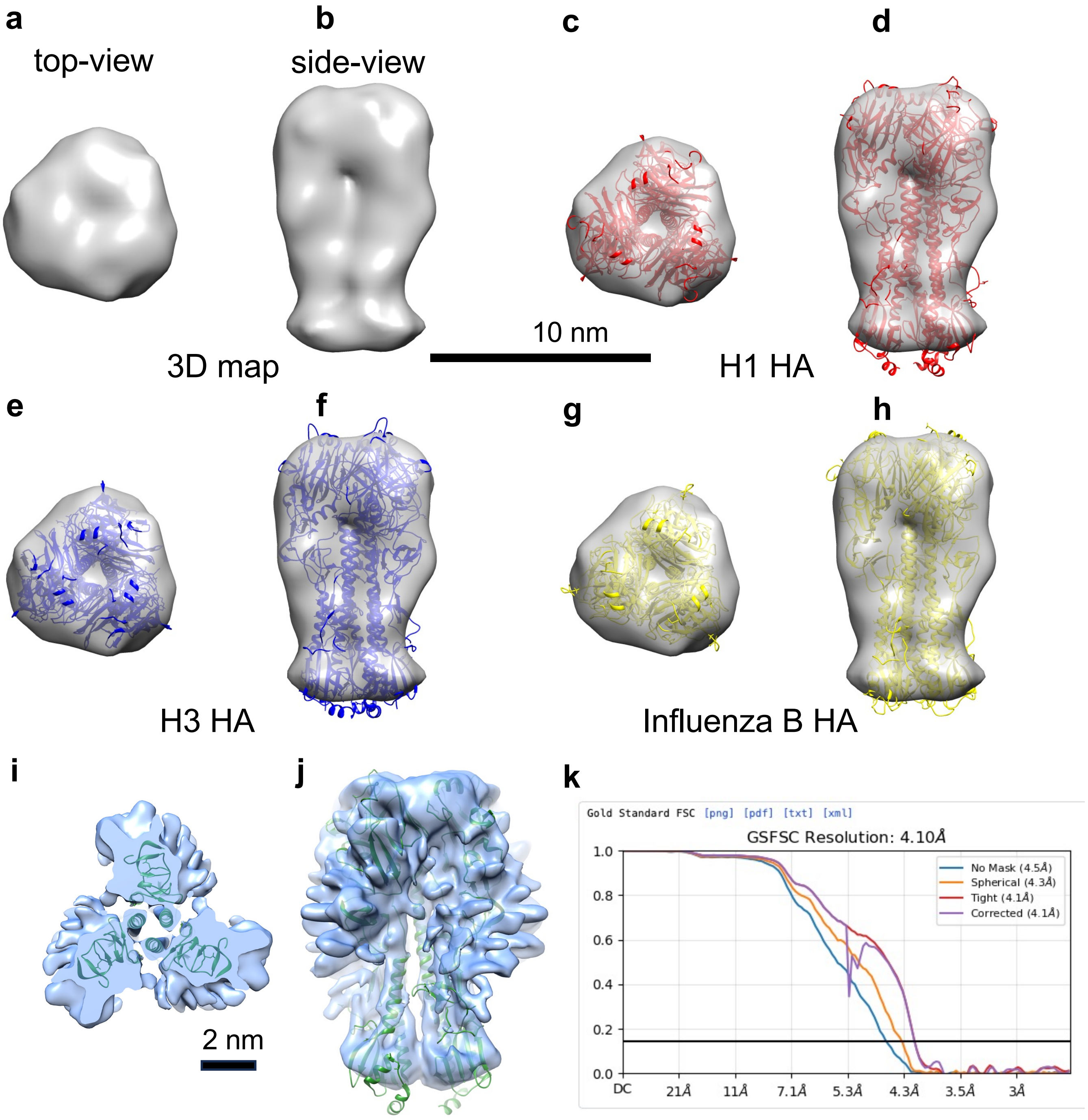


**Supplementary Fig. 9. Comparison of the docking of different HA coordinates into 3D maps of HA.** (a, b) Top and side views , respectively, of ab initio cryo-EM 3D reconstruction of HA spike. (c, d). Top and side views, respectively, of the docking of H1 HA ectodomain trimeric coordinates (PDB 7kna) into the 3D map (gray). (e, f). Top and side views, respectively, of the docking of H3 HA ectodomain trimeric coordinates (PDB 4fnk) into the 3D map (gray). (g, h). Top and side views, respectively, of the docking of influenza B HA ectodomain trimeric coordinates (PDB 2rfu) into the 3D map (gray). The 3D map is shown as an isosurface rendering (gray) and coordinates are ribbon diagrams with H1 (red), H3 (blue) and influenza B HA (yellow). Scale bar, 10nm. (i, j) Top-view (panel i) and side view (panel j) of the docking of H1 ectodomain coordinates (green ribbons) into a refined 3D map (light blue isosurface). Scale bar, 2nm. (k) Corresponding FSC curves (estimated resolution, 4.1 Angstroms).

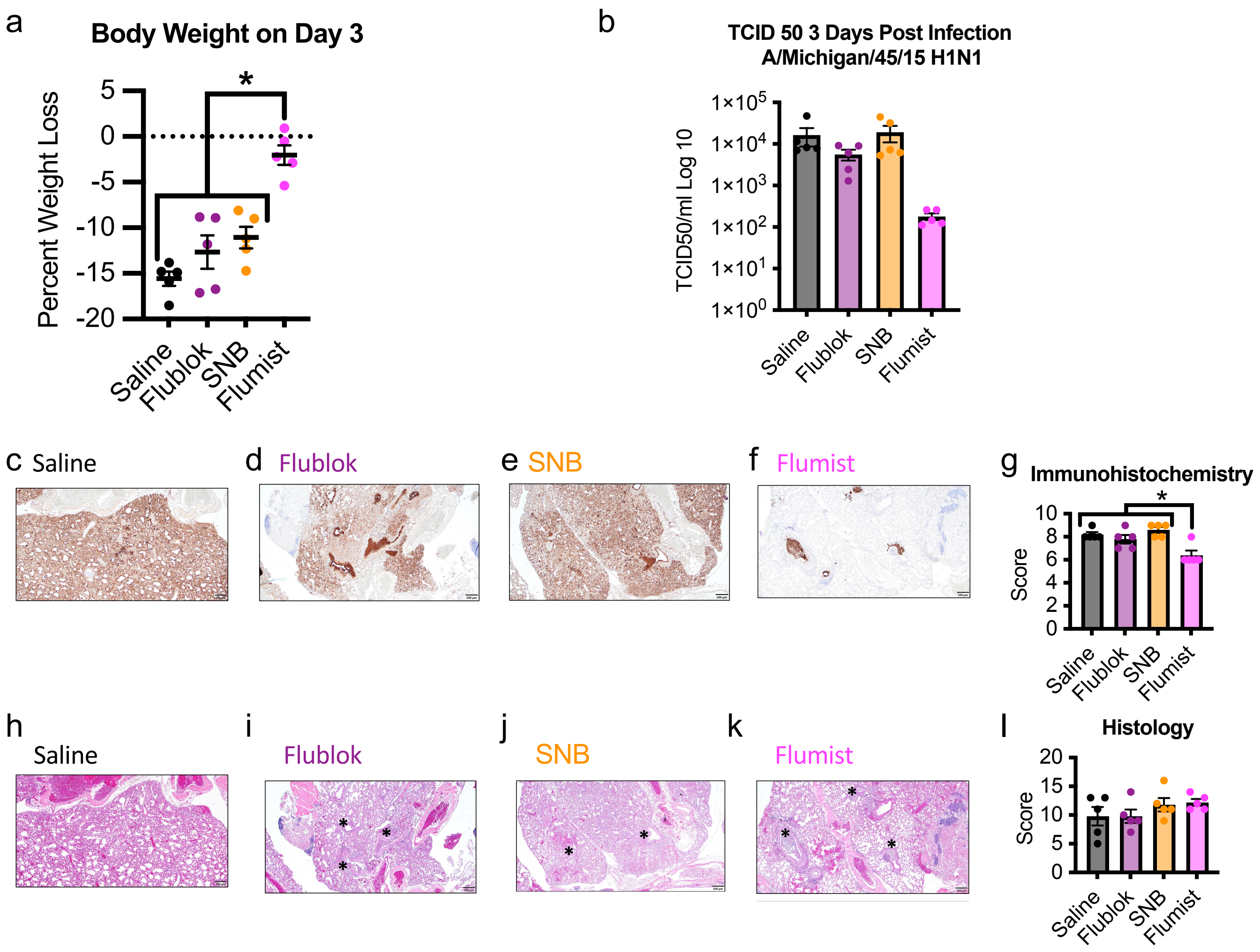


**Supplementary Fig. 10. Pathology comparisons of vaccines via intranasal immunization against H1N1 influenza virus (2015) challenge.** (a) Comparison of weight-losses on day 3. Graphs display mean with standard error of the mean bars. (b) Comparison of mouse lung titers determined by 50% Tissue Culture Infectious Dose (TCID50) for tissues taken day 3 post-H1N1 challenge with duplicate groups of mice. (c-f) Images show immunohistochemistry (IHC) against the influenza nucleoprotein protein for different vaccination groups and (g) scoring of images shown are from day 3 post H1N1 challenge for all groups. (h-k) Images of hematoxylin and eosin (H&E) staining of lung tissue sections for different vaccination groups and (l) histology scoring. Lung samples were collected from 2 mice per condition group and the experiment was replicated (N=4). All images at 10x magnification and display 100 μm scale bars. Immunization group colors are Saline (gray), Flublok (purple), Spiked nanobicelles (SNB) (orange) and Flumist (pink). Statistically significant differences are denoted with an asterisk to represent a significance level (p) less than 0.05.

**a**

ABO21709.1 HA A/PuertoRico/8/1934_H1N1
BCD56299.1 HA A/California/04/09 H1N1
ARH14956.1 HA A/Michigan/15/2015 H1N1

**b**

```
A/MI/25/15      MKAILVVLLYTFTTANADTLCIGYHANNSTDTVDTVLEKNVTVTHSVNLLEDKHNGKLCK  60
A/CA/04/09      MKAILVVLLYTFATANADTLCIGYHANNSTDTVDTVLEKNVTVTHSVNLLEDKHNGKLCK  60
A/PR/8/34       MKANLLVLLCALAAADADTICIGYHANNSTDTVDTVLEKNVTVTHSVNLLEDSHNGKLCR  60

A/MI/25/15      LRGVAPLHLGKCNIAGWILGNPECESLSTASSWSYIVETSNSDNGTCYPGDFINYEELRE  120
A/CA/04/09      LRGVAPLHLGKCNIAGWILGNPECESLSTASSWSYIVETPSSDNGTCYPGDFIDYEELRE  120
A/PR/8/34       LKGIAPLQLGKCNIAGWLLGNPECDPLLPVRSWSYIVETPNSENGICYPGDFIDYEELRE  120

A/MI/25/15      QLSSVSSFERFEIFPKTSSWPNHDSNKGVTAACPHAGAKSFYKNLIWLVKKGNSYPKLNQ  180
A/CA/04/09      QLSSVSSFERFEIFPKTSSWPNHESNKGVTAACPHAGANSFYKNLIWLVKKGNSYPKLSK  180
A/PR/8/34       QLSSVSSFERFEIFPKESSWPNHNTN-GVTAACSHEGKSSFYRNLLWLTEKEGSYPKLKN  179

A/MI/25/15      SYINDKGKEVLVLWGIHHPSTTADQQSLYQNADAYVFVGTSRYSKKFKPEIATRPKVRDQ  240
A/CA/04/09      SYINDKGKEVLVLWGIHHPPTSADQQSLYQNADTYVFVGSSRYSKKFKPEIAIRPKVRGQ  240
A/PR/8/34       SYVNKKGKEVLVLWGIHHPPNSKEQQNLYQNENAYVSVVTSNYNRRFTPEIAERPKVRDQ  239

A/MI/25/15      EGRMNYYWTLVEPGDKITFEATGNLVVPRYAFTMERNAGSGIIISDTPVHDCNTTCQTPE  300
A/CA/04/09      EGRMNYYWTLVEPGDKITFEATGNLVVPRYAFAMERNAGSGIIISDTPVHDCNTTCQTPK  300
A/PR/8/34       AGRMNYYWTLLKPGDTIIFEANGNLIAPMYAFALSRGFGSGIITSNASMHECNTKCQTPL  299

A/MI/25/15      GAINTSLPFQNIHPITIGKCPKYVKSTKLRLATGLRNVPSIQSRGLFGAIAGFIEGGWTG  360
A/CA/04/09      GAINTSLPFQNIHPITIGKCPKYVKSTKLRLATGLRNIPSIQSRGLFGAIAGFIEGGWTG  360
A/PR/8/34       GAINSSLPYQNIHPVTIGECPKYVRSAKLRMVTGLRNNPSIQSRGLFGAIAGFIEGGWTG  359

A/MI/25/15      MVDGWYGYHHQNEQGSGYAADLKSTQNAIDKITNKVNSVIEKMNTQFTAVGKEFNHLEKR  420
A/CA/04/09      MVDGWYGYHHQNEQGSGYAADLKSTQNAIDEITNKVNSVIEKMNTQFTAVGKEFNHLEKR  420
A/PR/8/34       MIDGWYGYHHQNEQGSGYAADQKSTQNAINGITNKVNTVIEKMNIQFTAVGKEFNKLEKR  419

A/MI/25/15      IENLNKKVDDGFLDIWTYNAELLVLLENERTLDYHDSNVKNLYEKVRNQLKNNAKEIGNG  480
A/CA/04/09      IENLNKKVDDGFLDIWTYNAELLVLLENERTLDYHDSNVKNLYEKVRSQLKNNAKEIGNG  480
A/PR/8/34       MENLNKKVDDGFLDIWTYNAELLVLLENERTLDFHDSNVKNLYEKVKSQLKNNAKEIGNG  479

A/MI/25/15      CFEFYHKCDNTCMESVKNGTYDYPKYSEEAKLNREKIDGVKLESTRIYQILAIYSTVASS  540
A/CA/04/09      CFEFYHKCDNTCMESVKNGTYDYPKYSEEAKLNREEIDGVKLESTRIYQILAIYSTVASS  540
A/PR/8/34       CFEFYHKCDNECMESVRNGTYDYPKYSEESKLNREKVDGVKLESMGIYQILAIYSTVASS  539

A/MI/25/15      LVLVVSLGAISFWMCSNGSLQCRICI  566
A/CA/04/09      LVLVVSLGAISFWMCSNGSLQCRICI  566
A/PR/8/34       LVLLVSLGAISFWMCSNGSLQCRICI  565
```

**c**

**Mutations in HA compared to A/Michigan/15/2015**

| | Full length HA0, 566 | | HA1, 20-344=324 | | HA2, 344-520=176 | | HA2/HA1 | |
|---|---|---|---|---|---|---|---|---|
| Strain | mutations | identity | mutations | identity | mutations | identity | ratio | % |
| A/California/04/2009 | 20 | 98% | 16 | 95% | 3 | 98% | 3/16 | 18% |
| A/PuertoRico/8/1934 | 108 | 81% | 81 | 75% | 15 | 91% | 15/81 | 18% |

**Supplementary Fig. 11. Sequence comparison of H1 HA proteins from 1934, 2009, and 2015 influenza viruses.** (a) GenBank accession numbers for the H1 HA protein sequences. (b) Sequence alignment of H1 HA sequences. Sequence regions corresponding the HA1 (red) and HA2 (blue) are denoted with amino acids differences colored (black). Arginine (R) is underlined marking the separation of HA1 and HA2 regions. (c) comparison of the number of mutations and sequence identity between A/Puerto Rico/8/1934 and A/Michigan/15/2015 H1N1 influenza viruses.

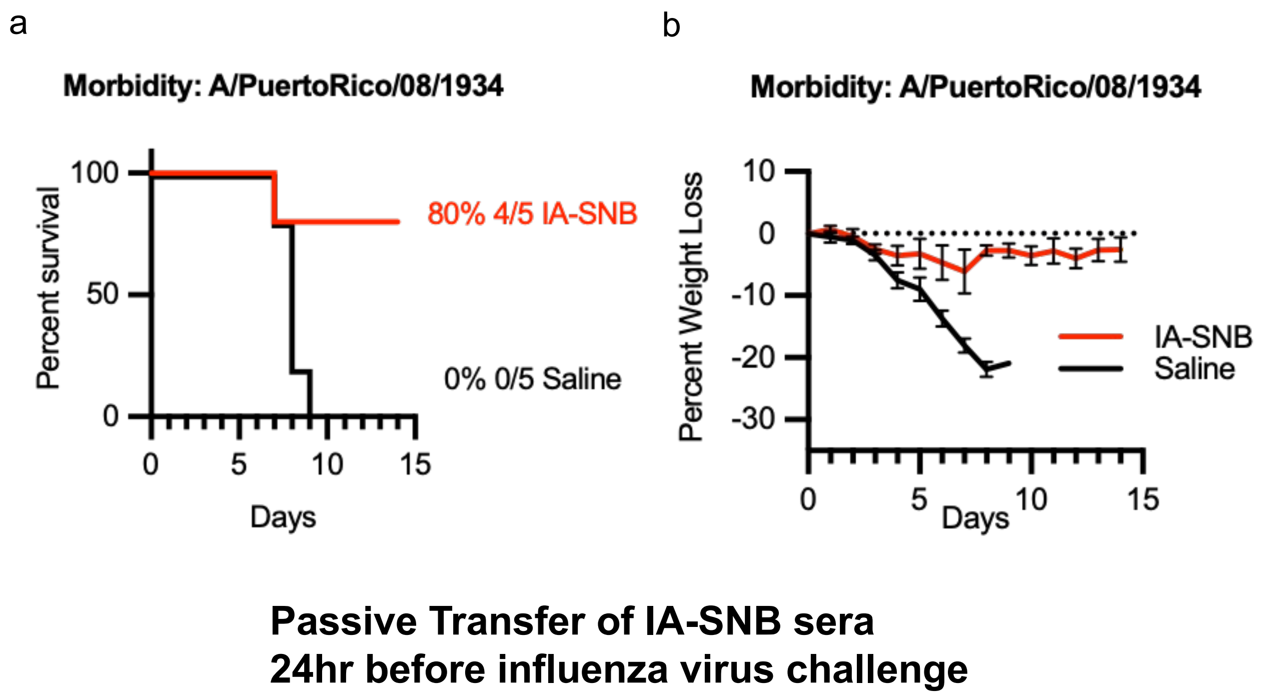


**Supplementary Fig. 12. Passive-transfer of vaccine sera of in vitro assembled spiked nanobicelles (IA-SNB) for protection from H1N1 challenge**. (a) Survival curves for mice that were intranasally challenged with influenza virus (A/Puerto Rico/8/1934 (H1N1)) 24 hours after intraperitoneally (IP) transfer of 400ul of saline sera (black line) or sera collected from mice immunized with 1934 H1 HA spiked nanobicelles (IA-SNB) (red line). (b) The corresponding weight-lost curves for panel a, with IA-SNB vaccine sera (red line) and saline negative-control (black line).

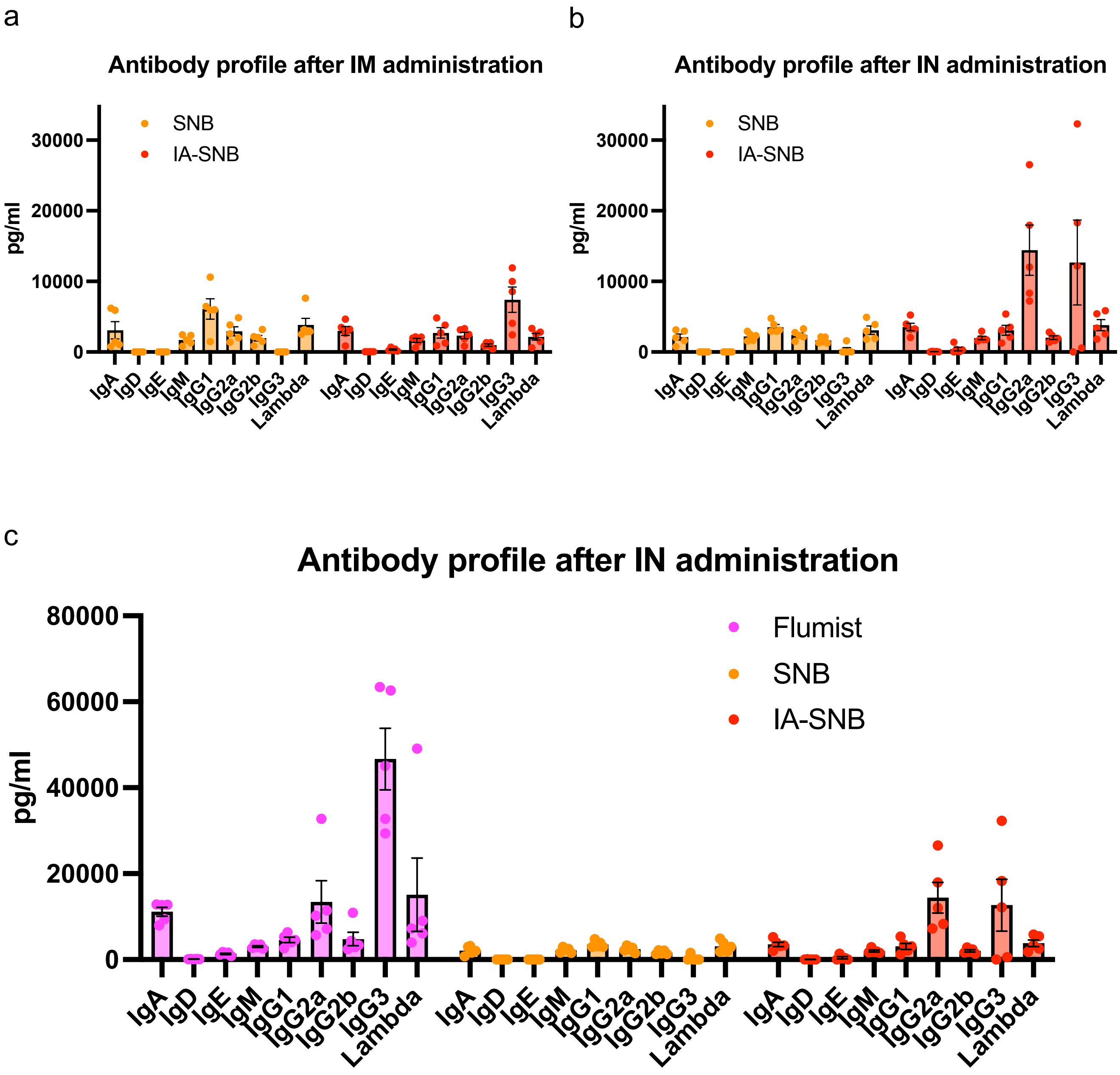


**Supplementary Fig. 13. Analysis of relative amount of different antibody classes and subclasses in different vaccine sera a day 35.** (a) Sera from mice intramuscularly immunized with spiked nanobicelles from Fluad (SNB)(orange) or in vitro assembled spiked nanobicelles (IA-SNB) (red) of 1934 H1 HA. (b) Sera from mice intranasally immunized with spiked nanobicelles from Fluad (SNB)(orange) and in vitro assembled spiked nanobicelles (IA-SNB) (red). (c) Comparing antibody classes and subclasses of sera from intranasally administered Flumist (pink) to intranasally administered Fluad SNB (orange) and IA-SNB (red).
